# A global resource constrained model to predict metabolic flux dynamics in fluctuating environments

**DOI:** 10.1101/2024.10.12.617971

**Authors:** Huili Yuan, Yang Bai, Xiongfei Fu

## Abstract

Environmental changes often induce global variations on bacterial gene expression, frequently accompanied by alterations in growth rates. Integrating non-metabolic cellular processes, such as gene expression and macromolecule synthesis, with metabolic modeling remains a significant challenge in systems biology due to the lack of mechanistic representation of gene regulation. Here, we introduce a novel constraint-based modeling framework called dynamic Constrained Allocation Flux Balance Analysis (dCAFBA), which comprehensively integrates metabolism, cellular resource allocation, and gene regulation. We employ a quasi-steady-state assumption, positing that reaction fluxes achieve balance at each time step, adapting more rapidly than protein synthesis and growth dilution. This approach enables the prediction of reaction flux dynamics and protein allocation necessary to achieve cellular objectives, such as optimizing growth during various growth shifts (including carbon, amino acid, and transcriptional signal), without detailing molecular gene regulations. Our model offers a new method for interpreting proteome allocation and cellular metabolism in complex and transient environments, providing mechanistic insights valuable for metabolic engineering.

## Introduction

Living organisms frequently encounter fluctuating environmental conditions. For instance, *Escherichia coli* in the mammalian intestine faces significant variability in nutrient availability, such as carbon and nitrogen sources, at different intestinal locations and different times of a day. Additionally, microbial cells experience genetic variations, either naturally or experimentally, such as through the induction of specific genes by controlling regulatory signals. To cope with these changes, bacteria must timely reshape global gene expression and redistribute metabolic flux to meet its metabolic requirements and protein synthesis (Figure 1). Investigating the kinetic responses of gene expression and metabolic distribution to changes in growth conditions is crucial for understanding adaptive strategies in microorganisms. While gene expression responses have been extensively studied, ^1-5^ the metabolic distribution responses remain less reported.

**Figure 1.**
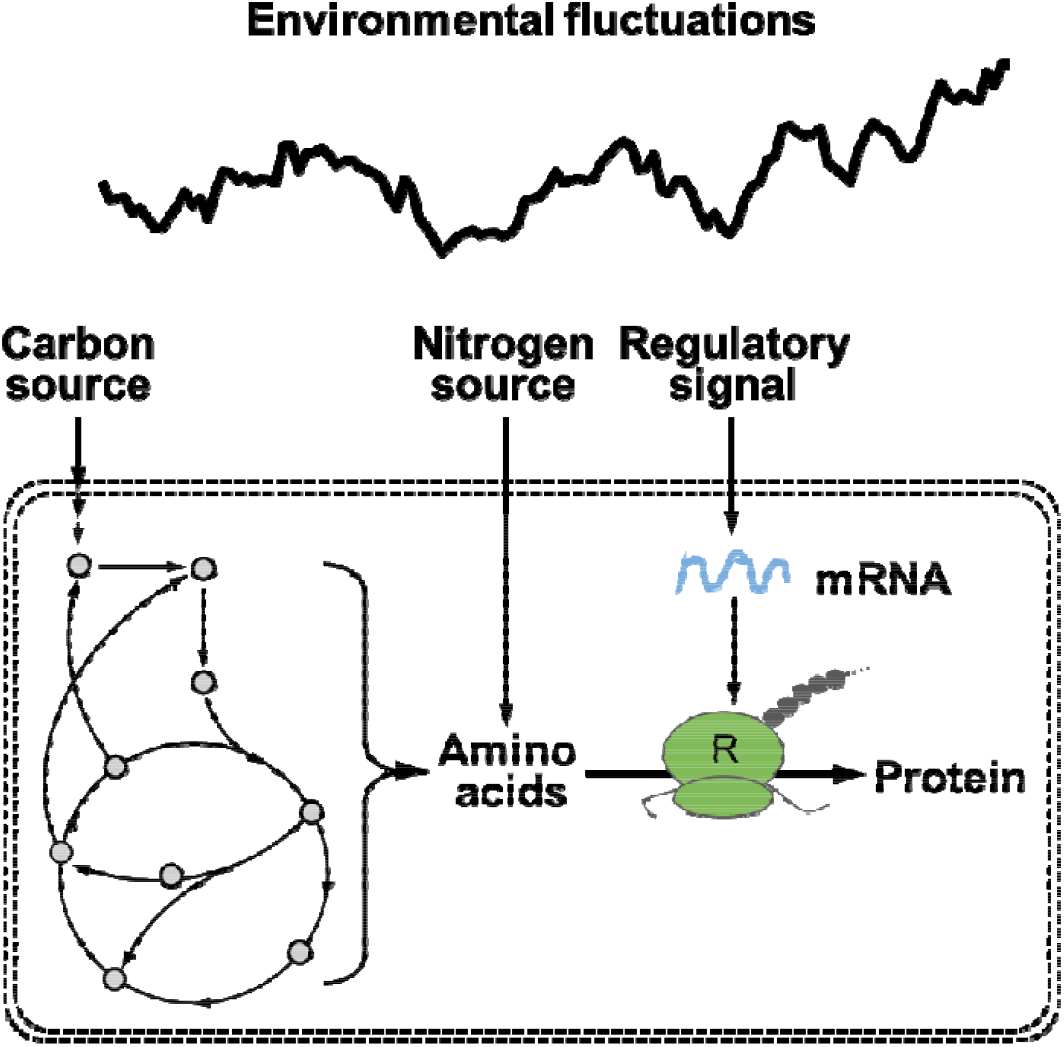
The kinetic responses of metabolic fluxes and protein synthesis to environmental fluctuations, including time-varying carbon source, nitrogen source and regulatory signal.

Constraint-based modelling, such as flux balance analysis (FBA), has been proven a powerful tool for predicting and analyzing metabolic flux distributions using genome-scale metabolic models (GSMMs). ^6-8^ However, traditional FBA assumes a steady-state condition for intracellular metabolites, which limits its ability to simulate cellular transient behaviors under fluctuating conditions, such as nutrient availability and nutrient shifts. To address this, dynamic FBA (dFBA) was developed, solving the FBA problem iteratively at each time step by updating the extracellular nutrients accordingly. ^9^ However, like classic FBA, dFBA only employs stoichiometric constraints through the mass conservation of metabolites, ignoring the building costs of proteins and molecular machines such as ribosome. To improve the accuracy of dFBA, several frameworks have been developed to integrate additional biological information, such as protein constraints. ^10-12^

Despite improved predictions and applications, constructing protein-constrained metabolic models remains challenging. One major issue is the dependence on enzymatic parameters (*k*_*cat*_) to further constrain the solution space of possible flux distribution. These essential parameters are often not readily available at the genome-scale, and are obtained from non-physiological conditions. ^13^ One approach to solve this issue is by incorporating phenomenological growth laws into FBA, which avoids the requirement for detailed enzymatic kinetic data. ^14,15^ Proteome allocation, the basis of bacterial growth laws and coarse-grained mechanistic models, plays a central role in relating gene expression to cellular metabolism and growth rates under different growth conditions. ^16-20^ Protein resource allocation has also been suggested to govern growth transitional kinetics during nutrient shifts in a global manner. ^2-4^

In this work, we developed a dynamic modelling method called dynamic constrained allocation FBA (dCAFBA). By integrating the flux-controlled regulation model (i.e., FCR) ^2^ with metabolic models, we studied the growth transition kinetics and metabolic flux reconfigurations during environmental changes, including carbon shifts, amino acid (AA) shifts and perturbations in transcriptional signals. Our model provides a novel method to predict the growth transition kinetics and metabolic flux configurations under complex and transient environments, and provides mechanistic guidance for metabolic engineering.

## Results

### Construction of a dynamic metabolic model constrained by proteome allocation for various changing conditions

Microbes must redistribute their metabolic fluxes during environmental fluctuations to facilitate the synthesis of amino acids and other intracellular metabolites. These metabolites serve as precursors for protein biosynthesis, influencing the rate of protein production () and the translational activity of ribosomes (). The translational activity is a key parameter that governs the proteins allocation among various functional sectors, including catabolic proteins (C sector), ribosomal proteins (R sector), house-keeping proteins (Q sector) and metabolic enzymes (E sector). ^2,21^ Changes in protein allocation impact metabolic flux redistribution by imposing constraints on the maximal fluxes that metabolic reactions could carry.

To dynamically integrate flux redistribution and proteome allocation, we first analyzed the typical time scales of both processes. The transition time for metabolites to reach a balanced state is determined by their concentrations and metabolic reaction fluxes. High fluxes and low metabolite concentrations of central carbon metabolism enable substrates to rebalance within approximately 10 seconds. ^22^ In contrast, the timescale for proteome reallocation is significantly slower (on the order of 10-100 minutes) due to the slow synthesis and degradation of proteins, which are related to growth rates. This difference in time scales (over two orders of magnitudes) allows us to assume that metabolite concentrations quickly reach a steady state during changes in protein allocation.

This quasi-steady-state assumption enabled us to use flux balance analysis (FBA) to calculate the distribution of metabolic fluxes by optimizing the overall biomass synthesis rate, ^6^ while protein allocations *ϕ* were determined dynamically. Under fluctuating environments, the gene expression level of each protein sectors (characterizing by the regulation function *χ*(*σ*)) may change promptly with *σ* through rapid signaling responses mediated by cAMP and ppGpp. ^17^ However, the allocation of each protein sectors *ϕ* must follow the slower dynamics of protein accumulation and dilution, governed by the protein synthesis rate *ν*_*R*_(*t*) (unit in per mass) and the dilution rate (i.e., growth rate *λ*):

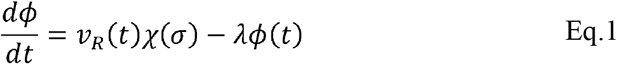

By fitting the regulation function *χ*(*σ*) from proteomics data of steady-state gene expression levels at various growth rates (see Methods), we derived the dynamics of *ϕ*(*t*). These protein allocations then constrained the upper bounds of metabolic fluxes in FBA to determine the dynamics of metabolic fluxes (see Methods).

The remaining component of the model is the determination of the translational activity *σ*(*t*), defined by the protein synthesis rate *ν*_*R*_(*t*) and ribosome allocation *ϕ*_*R*_(*t*), i.e., 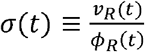. In most cases where all the amino acids are produced or transported coordinately, the protein synthesis rate *ν*_*R*_(*t*) can be considered proportional to the global amino acid synthesis rate (*ν*_*aa*_(*t*)) calculated from FBA: *ν*_*R*_(*t*) ∝ *ν*_*aa*_(*t*). The dynamics of translational activity can be derived by its full derivative:

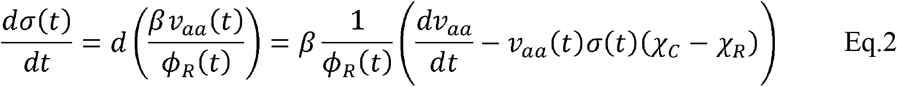

In the case of amino acids depletion, the biosynthesis of individual amino acid is turned on in order. The dynamics of *σ*(*t*) should be represented discontinuously using stepwise functions, which will be detailed in the section of amino acid down-shift.

To summarize, we constructed the global resource allocation model by categorizing all proteins into four sections based on their response to translation activity *σ*(*t*). The dynamics of each coarse-grained protein allocations *ϕ*_*i*_(*t*) were then calculated using the protein synthesis flux *ν*_*R*_(*t*) and regulation functions *χ*_*i*_(*σ*) determined experimentally. Applying the quasi-steady-state assumption to metabolic fluxes and constraining these fluxes by the protein allocations *ϕ*_*i*_(*t*), we predicted the distribution of metabolic fluxes using FBA methods. The calculated amino acid synthesis flux *ν*_*aa*_(*t*) then determined the protein synthesis fluxes *ν*_*R*_(*t*) and the translational activity *σ*(*t*) either continuously or discretely depending on the context.

### Kinetic response of metabolic fluxes during carbon shifts

To understand the regulatory functions during carbon shifts, where growing cells transition from a single carbon substrate to two co-utilized carbon substrates or vice versa, we described the protein synthesis of the C and R sectors using the FCR model. ^2^ Specifically, the regulation functions of protein allocations *χ*_*C*_ (*σ*) and *χ*_*R*_ (*σ*) are described as 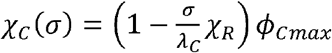 and 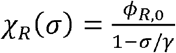, where *λ*_*C*_, *γ, ϕ*_*Cmax*_, *ϕ*_*R*,0_ are constants determined by steady-state protein allocations under different growth conditions. ^2^ Using these regulation function *χ*_*C*_ and *χ*_*R*_, the dynamics of each protein sector can be described by 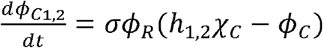 and 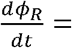 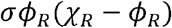. The parameter *h*_1,2_ denotes the partition of a carbon substrate in a pair of co-utilized carbon sources and is fitted to ensure the predicted growth rate matches experimental data.

Our dCAFBA model, which integrated the FCR model with FBA, captured both proteome allocation changes and the kinetics of metabolic fluxes during growth transitions. By comparing with the experimental data, we found that dCAFBA successfully captured the transition kinetics of growth rate *λ* during both carbon up-shifts and carbon down-shift (Figure S1). During carbon up-shifts, the growth rate gradually increased from a lower value to a higher steady-state value. Conversely, upon the depletion of a second carbon source in a carbon down-shift, the cellular growth rate dropped abruptly and then gradually increased until a new steady-state was reached. Notably, an overshoot in the growth rate was observed during the carbon down-shift, resulting from limitation switch from carbon uptake proteins to metabolic enzymes. ^21^ We also presented the dynamics of carbon uptake proteins *ϕ*_*C*_, ribosomal proteins *ϕ*_*R*_ and their regulatory functions *χ*_*C*_, *χ*_*R*_ during carbon shifts in Figure S1.

Importantly, the dCAFBA model can predict the kinetics of metabolic fluxes during growth transitions. We first investigated the dynamics of the metabolic reactions in the central carbon metabolic pathways during a carbon up-shift from succinate to a combination of succinate and glycerol. The co-utilization of succinate and glycerol resulted in a higher growth rate compared to single substrates. The glycerol uptake reaction (denoted as ‘GLYK’) was activated immediately after the shift, with its flux gradually increasing (Figure 2b). This increase is due to the expression of proteins related to glycerol uptake and transport upon the presence of glycerol. Subsequently, the glycerol influx was directed into reactions catalyzed by glycerol phosphate dehydrogenase (‘G3PD2’) and triose phosphate isomerase (‘TPI’), which exhibited similar flux kinetics to glycerol uptake. Upon the addition of glycerol to the medium, the succinate uptake flux (‘Succ_ext_’) gradually decreased, leading to similar flux patterns in downstream reactions such as succinate dehydrogenase (‘SUCD1i’) and fumarase (‘FUM’). This reduction is expected as the expression of the succinate uptake system is significantly reduced in the presence of glycerol. ^23^ Interestingly, we observed that fluxes going into reactions catalyzed by fructose bisphosphatase (‘FBP’) and fructose bisphosphatase aldolase (‘FBA’) dropped to zero at the shift time. This is due to the feedback inhibition by the glycolytic intermediate fructose-1,6-biphosphate (FBP), which suppresses glycerol uptake. ^24^

**Figure 2.**
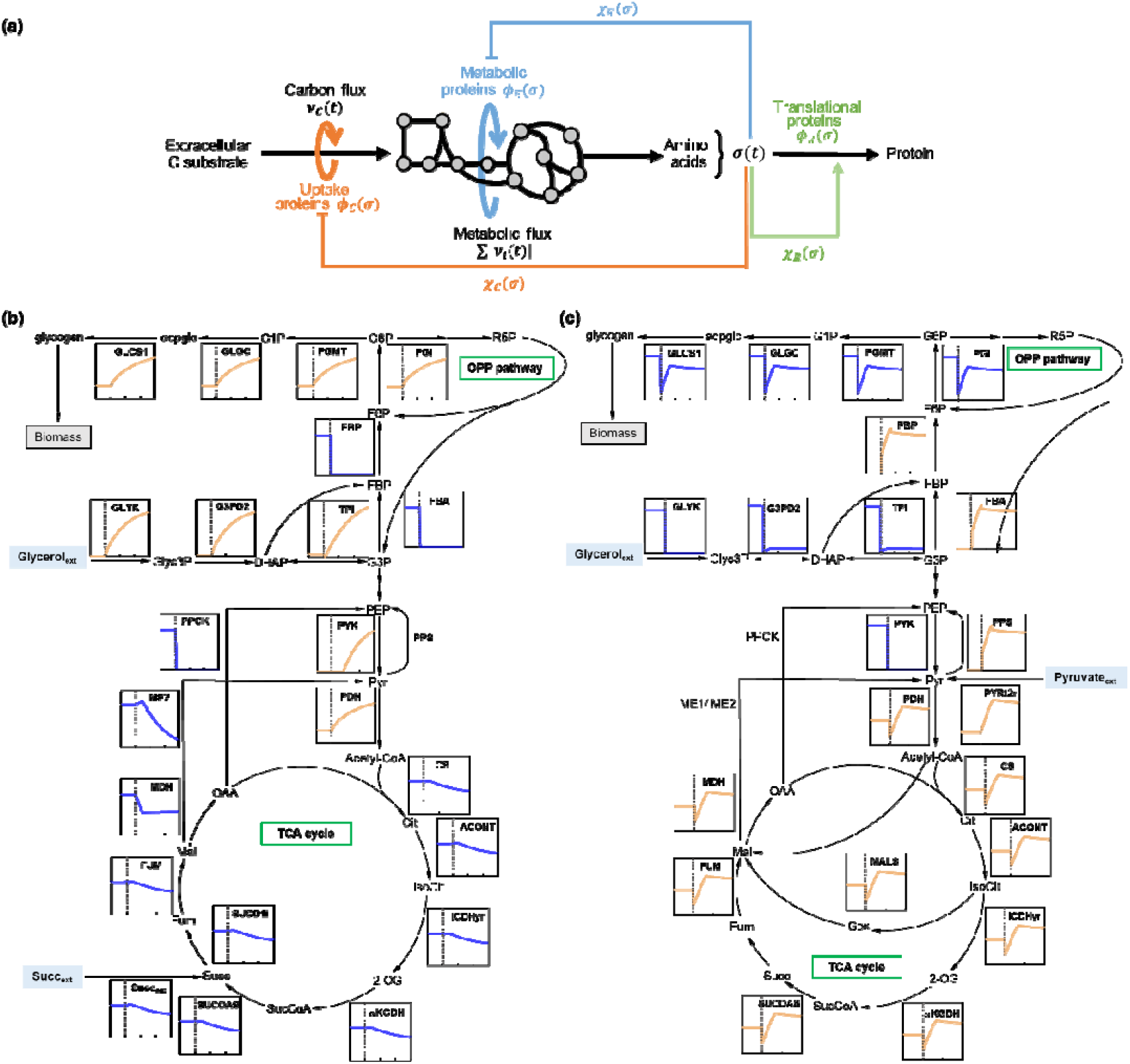
Model illustration of carbon shift and simulated results. (a) A schematic illustrating the integration of protein synthesis within the *E*.*coli* metabolic network, incorporating regulation functions of protein synthesis and flux constraints. This integration allows for the prediction of protein allocation and metabolic fluxes. Extracellular carbon substrates are taken in by uptake proteins in the carbon influx and are converted into central precursors and amino acids by metabolic proteins. These amino acids, which affect the ribosome’s translational activity, are further consumed by the ribosome for synthesizing uptake (), metabolic (), translational () proteins, and others. The regulatory effects are implemented through regulation functions,, and, which determine the amounts of uptake, metabolic and ribosomal proteins, respectively. (b) The flux dynamics of selected reactions in the glycolysis, gluconeogenesis and TCA pathways during the transition from succinate to a combination of succinate and glycerol. The x-axis represents time (in hours), and the y-axis represents reaction flux (mmol/gDW/h). The vertical grey line at 0 hour indicates the shifting time, separating the pre-shift (left) from the post-shift. (c) The flux dynamics of the selected reactions in the glycolysis, gluconeogenesis and TCA pathways during the transition from pyruvate and glycerol to pyruvate. Abbreviations: Glycerol_ext_, extracellular glycerol; G1P, glucose-1-phosphate; G6P, glucose-6-phosphate; F6P, fructose-6-phosphate; FBP, fructose-1,6-biphosphate; G3P, glyceraldehyde-3-Phosphate; DHAP, dihydroxyacetone phosphate; PEP, phosphoenol pyruvate; Pyr, pyruvate; OAA, oxalacetic acid; Cit, citrate; IsoCit, threo-isocitrate; Succ, succinate; Succ_ext_, extracellular succinate; Pyruvate_ext_, extracellular pyruvate; SucCoA, succinyl-CoA; Fum, fumarate; Mal, malate; 2-OG, 2-oxoglutarate; adpglc, ADP-glucose; Gly3P, glycerol-3-phosphate; PGI, glucose-6-phosphate isomerase; PFK, phosphofructokinase; FBP, fructose bisphosphatase; FBA, fructose bisphosphate aldolase; TPI, triose phosphate isomerase; PPS, phosphoenolpyruvate synthase; PYK, pyruvate kinase; CS, citrate synthase; ACONT, aconitase; ICDHyr, isocitrate dehydrogenase; AKGDH, 2-Oxogluterate dehydrogenase; SUCOAS, succinyl CoA synthetase; SUCD1i, succinate dehydrogenase; FUM, fumarase; MDH, malate dehydrogenase; PPCK, phosphoenolpyruvate carboxykinase; ME1/ME2, malic enzyme; PGMT, Phosphoglucomutase; GLGC, glucose-1-phosphate adenylyltransferase; GLCS1, glycogen synthase; G3PD2, glycerol-3-phosphate dehydrogenase.

In contrast to the carbon up-shifts, the changes in metabolic fluxes during the carbon down-shifts were distinct. During the co-utilization of pyruvate and glycerol, both substrates were simultaneously consumed in the pre-shift, represented by ‘PYRt2r’ and ‘GLYK’, respectively (Figure 2c). In the absence of glycerol, the reaction catalyzed by glycerol kinase (‘GLYK’) was turned off post-shift. However, the flux of ‘G3PD2’ remained nonzero in the post-shift as ‘G3PD2’ operated in the reverse direction, producing glycerol-3-phosphate (Glyc3P) from dihydroxyacetone phosphate (DHAP). In the post-shift, the pyruvate uptake reaction remained active, exhibiting two distinct transition stages. Initially, the pyruvate uptake flux increased rapidly, accompanied by an increasing in the expression of its uptake proteins. In the second stage, the pyruvate influx gradually decreased until a final steady state was achieved. Similarly, the flux dynamics of other reactions, such as pyruvate dehydrogenase (PDH) and citrate synthase, also exhibited an overshoot phenomenon, which has been extensively discussed in our recent work. ^21^ In addition to studying the *E*.*coli i*JR904 metabolic network ^25^, we also applied our dCAFBA model to the latest metabolic model of *E*.*coli, i*ML1515, ^26^ following the same grouping of reactions as in *i*JR904 metabolic model. The resulting simulations were similar to those obtained using *i*JR904 (Figure S2).

### Sequential express of AAB enzymes and redistribution of metabolic fluxes during amino acid down-shift

To simulate the metabolic flux changes during an amino acid (AA) down-shift (transition from rich medium to amino acid-free minimal medium), we incorporated elements of the two-stage allocation model proposed by Wu et al. ^2^ into dCAFBA framework (Figure 3a). Upon AA removal, cells need to synthesize AAB enzymes (indicated by protein fractions *ϕ*_*A*1_, *ϕ*_*A*2_, etc.) to supply the missing AAs. The synthesized AAs, consumed by ribosomes for protein synthesis, reflect the ribosomal activity *σ*(*t*), defined as 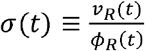. Following the FCR scheme ^2^, the translational activity *σ*(*t*) controls the allocation of protein synthesis flux to total AAB enzymes and ribosomal proteins via the regulatory functions *χ*_*A,tot*_ and *χ*_*R*_, respectively. These functions are mathematically expressed as 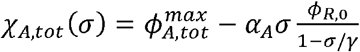 and 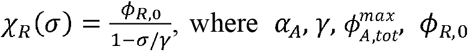 are constants determined from steady-state measurements pre-shift and post-shift.

**Figure 3.**
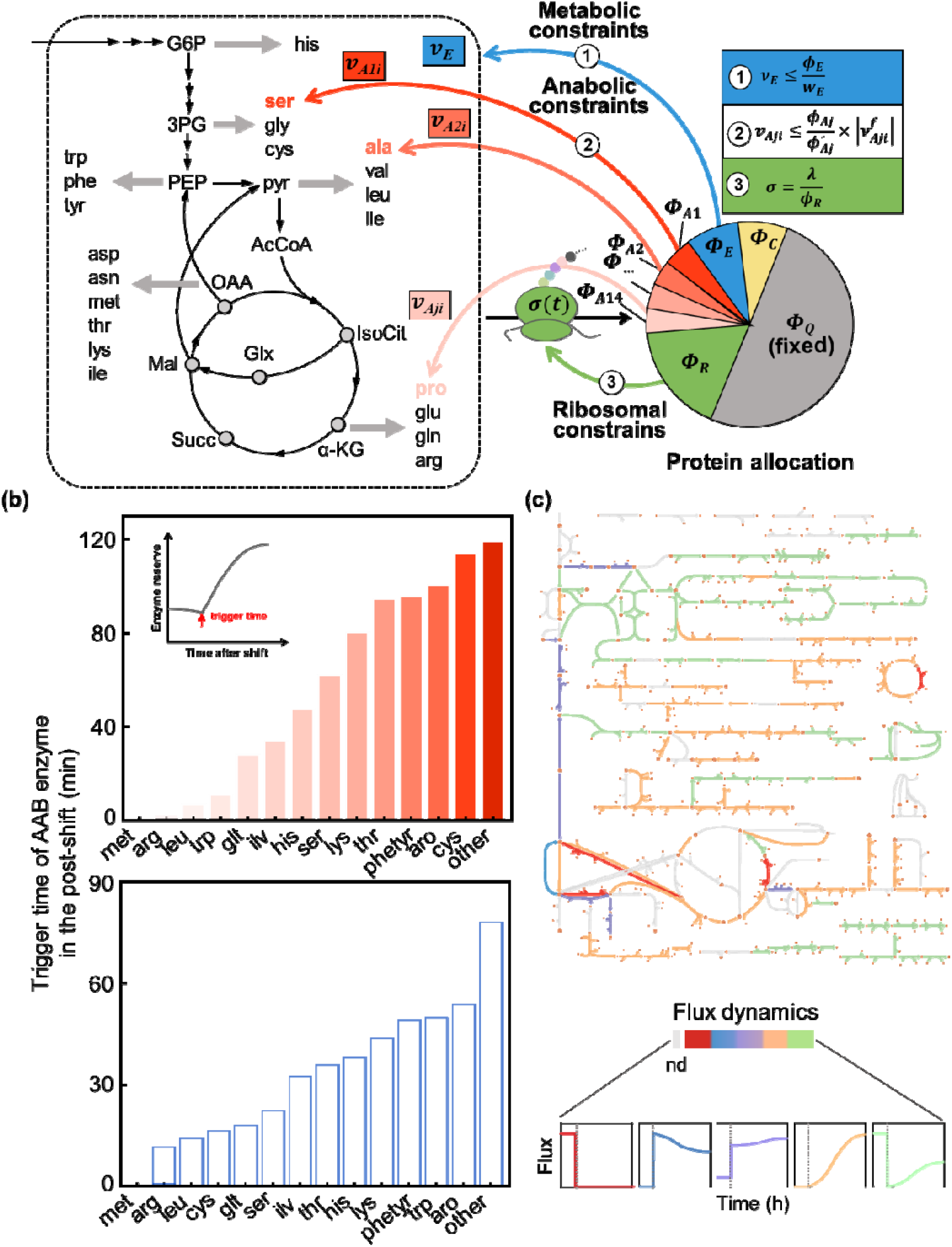
Model illustration of AA down-shift and its simulated results. (a) The taken-in substrates are converted to amino acids through a series of metabolic reactions,with fluxes constrained by metabolic proteins. The pool of amino acids reflects the ribosomal translational activity, which controls the allocation of protein synthesis flux to catabolic proteins (, AAB enzymes (, metabolic proteins (, ribosomal proteins (and house-keeping proteins (. The reaction flux of each AAB pathway is constrained by its protein abundance level (relative to the final post-shift level (as well as its flux under the steady state in the post-shift 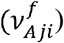. The model dynamics are set by the translational activity *σ*(*t*), with metabolic fluxes assumed to be in quassi-steady state. (b) The trigger time, defined as the time each AAB enzymes initiates expression in the post-shift. Protein groups are sorted in order of increasing trigger time. Upper panel: shift from glycerol with 18AAs to glycerol. Lower panel: shift from glucose with cAA to glucose. (c) Illustration of changes in reaction fluxes during AA down-shift. Colors represent the flux dynamics. This figure was created using Escher. ^27^

Based on end-product inhibition, the protein synthesis flux directed to total AAB enzymes *χ*_*A,tot*_ is further sub-allocated to the enzymes of individual pathways (for example, alanine, leucine) by the pathway-specific allocation function *χ*_*A,j*_(*t*) ≡ *η*_*j*_(*t*)*χ*_*A,tot*_. The group-specific relative allocation functions *χ*_*A,j*_(*t*)are expressed as

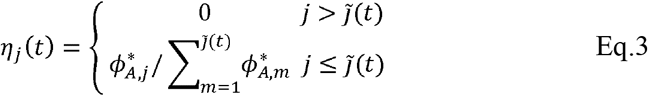

where 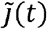 is the number of enzyme groups being synthesized at the time *t*. Groups with 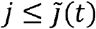 are growth-limiting, while those with 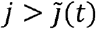 are not. Whether an enzyme group is growth-limiting is determined by its present enzyme abundance *ϕ*_*Aj*_ relative to the final abundance 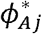 denoted as 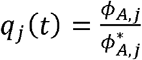. At any time t, the fractional enzyme reserve *q*_*j*_(*t*) divides the AAB groups into two classes: non-limiting groups receive no allocation (*η*_*j*_(*t*) =0), and limiting groups with the lowest *q*_*j*_(*t*) are allocated in proportion to their final steady state abundance 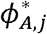. Referring to Wu et al. ^3^, the total AAB enzymes are classified into 14 groups based on operon structure and 89 reactions (see Table S5), with the sum of the fourteen *χ*_*A,j*_ equaling *χ*_*A,tot*_.

We first examined the kinetics of the growth rate *λ*, proteome fractions (*ϕ* _*R*_, *ϕ*_*A,tot*_) and their regulatory functions (*χ*_*R*_, *χ*_*A,tot*_) during the AA-shift. During the transition from glycerol supplemented with 18AAs (Table S10) to glycerol alone (Figure S3a), our predicted growth kinetics were consistent with experimental results. In the post-shift, the regulatory function for ribosomal proteins *χ*_*R*_ was repressed, while the regulatory function for total AAB enzymes *χ*_*A,tot*_ was upregulated. The dynamics of proteome fractions *ϕ*_*R*_ and *ϕ*_*A,tot*_ were predicted to align well with experimental data.

As described in the model construction section, the protein synthesis flux directed to total AAB enzymes is further sub-allocated to individual AAB groups according to the fractional enzyme reserve *q*_*j*_(*t*). We assumed cells preferentially allocate protein synthesis flux to enzyme groups with the smallest protein reserve. Using this assumption, we computed the protein reserve *q*_*j*_(*t*) for each AAB enzyme after the shift (Figure S4a). We observed that met group was synthesized immediately after the shift because methionine, with the smallest reserve at the shifting time (0 hour, *q*_*j*_(0)), was the most bottlenecked. Once cellular growth was no longer limited by the met group, other AAB enzyme groups began to express sequentially, leading to a stepwise expression of individual AAB enzymes groups. We defined the time at which each AAB enzyme group initiate to express s in the post-shift as the trigger time, essentially determined by its protein reserve *q*_*j*_(*t*). We then sorted the 14 individual AAB groups in order of increasing trigger time (Figure 3b). The predicted results closely matched experimental observations. ^3^ Using the calculated protein reserve *q*_*j*_(*t*), we obtained the sub-allocation regulatory function for each AAB group *χ*_*A,j*_ (Figure S4b).

We repeated the AA down-shift simulations for other conditions, including the transition from glycerol supplemented with 18AAs without serine to glycerol (Figure S3b), and the transition from glucose supplemented by casamino acid (cAA) to glucose minimal medium (Figure S5), obtaining very similar results. We also computed the trigger time of AAB enzymes for each condition, which varied as expected due to differences in the steady-state protein fractions of each AAB enzyme group under different growth conditions. However, in all the three cases, the met group was the first to be activated after the shift. In addition to studying the *E*.*coli i*JR904 metabolic network, we tested the AA down-shift model on the latest *E*.*coli* metabolic reconstruction, *i*ML1515, ^26^ generating very similar results (Figure S6). These analyses suggested our model was robust across different growth media and metabolic models.

Our dCAFBA model enabled us to capture the time courses of metabolic flux during AA down-shift. We expected significant changes in metabolic flux due to the requirement for *de novo* AA biosynthesis to maintain cellular growth in the absence of external AAs. Our predicted results showed diverse responses in metabolic flux changes through specific reactions (Figure 3c). Fluxes of some reactions dropped to zero immediately after the shift, such as those catalyzed by pyruvate dehydrogenase (PDH) and phosphoenolpyruvate carboxykinase (PPCK). In contrast, other reactions, such as phosphoenolpyruvate synthase (PPS), exhibited fluxes that abruptly increased at the shift time (0 hour) and then gradually decreased.

Based on the patterns of reaction fluxes, we divided the reactions in the central metabolic pathway into five groups, each assigned a color for visualization (Figure 3c). The results suggested that fluxes carried by lower glycolytic reactions (Figure 3c, purple lines), such as phosphoglycerate kinase (PGK) and enolase (ENO), increased immediately after the shift and then gradually increased post-shift. In contrast, most reactions involved in amino acid biosynthetic pathways (Figure 3c, orange lines) began increasing their fluxes post-shift until reaching a new steady-state value, such as tyrosine transaminase and isoleucine transaminase. This gradual increase is expected, as amino acids need to be synthesized in the post-shift medium lacking AAs. The cells prioritize the synthesis of these essential building blocks to support continued growth and protein production.

Overall, our model provides a comprehensive view of how metabolic fluxes are redistributed during AA down-shift, highlighting the dynamic adjustments cells make to maintain growth and function under nutrient-limited conditions. These insights are valuable for understanding bacterial adaptation and can inform strategies for metabolic engineering.

### Cellular dynamics under transcriptional variations

To further demonstrate the application of the developed dCAFBA model, we explored the growth dynamics perturbed by the inducible exogenous genes expression for valuable metabolites, such as pol-3-hydroxybutyrate (PHB), which is considered as a potential substitute for traditional plastics. ^28^ For this purpose, the *E*.*coli i*JR904 genome-scale metabolic model was modified by adding the PHB biosynthetic reactions (HACD1, 3HBC3E, PHB_syn) encoded by *phaA* (3-ketothiolase), *phaB* (acetoacetyl-CoA reductase), and *phaC* (PHB synthase), respectively. ^29^ These newly added reactions were assigned to a new *P* sector, and PHB synthesis proteins were denoted as *ϕ*_*P*_. Considering the additional flux drain included in the metabolic model, exogenous and endogenous enzymes would compete for the shared precursors acetyl-CoA. We assumed that the fluxes entering the PHB biosynthetic pathway and other metabolic reactions are allocated linearly with the ratio of their relative protein sectors, i.e. 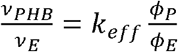. The parameter *k*_*eff*_ represents the competition for shared precursors between PHB synthesis and other metabolic reactions in the metabolic network (Figure 4a).

**Figure 4.**
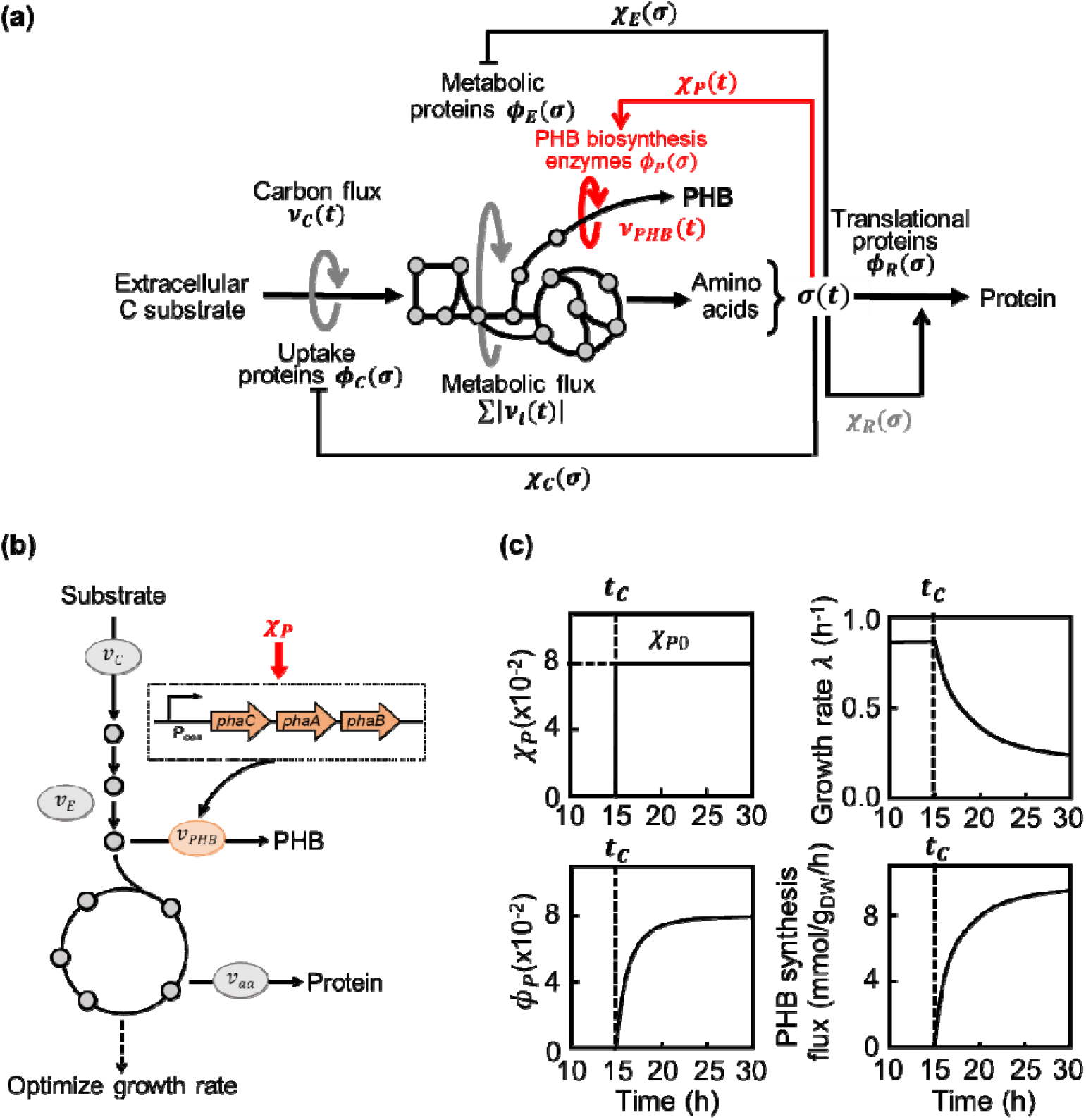
Model illustration of exogenous gene expression and simulated results. (a) Schematic illustration of the dCAFBA model simulating growth dynamics perturbed by enhanced PHB production. Extracellular carbon substrates are taken in by uptake proteins in the carbon influx and are converted into carbon precursors and amino acids by metabolic proteins. These amino acids are consumed by the ribosome for synthesizing uptake (), metabolic (), translational () proteins, PHB biosynthesis-related enzymes (), and others. The regulatory functions and control the protein synthesis flux allocated to uptake and ribosomal proteins, respectively. (b) The PHB production module includes *phaA* (3-ketothiolase), *phaB* (acetoacetyl-CoA reductase), and *phaC* (PHB synthase). The flux of PHB synthesis is assumed to be determined by its protein fraction relative to the metabolic protein fraction and the total metabolic flux. The protein related to PHB synthesis () is regulated by a step function. (c) Regulation function of PHB synthesis protein, protein fraction of PHB synthesis, growth rate, and PHB synthesis upon inducing exogenous genes at and.

In the extended *E*.*coli* model, a step function *χ*_*P*_ controlling the PHB protein synthesis flux was introduced:

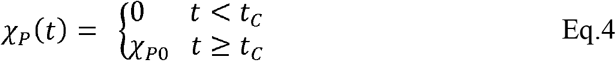

to present the induction of PHB synthesis gene expression at time *t*_*C*_ to level *χ*_*P*0_.

Similar to the *ϕ*_*C*_(*t*) and *ϕ*_*R*_(*t*), the protein fraction related to the PHB biosynthesis is also regulated by the translational activity *σ*(*t*) in the form: 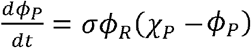. Once *ϕ*_*L*_(*t*_*n*_) and *ϕ*_*E*_(*t*_*n*_) at each time step *t*_*n*_ have been determined, the PHB synthesis flux *ν*_*PHB*_(*t*_*n*_) can be calculated using the total metabolic flux from the previous time step *ν*_*E*_(*t*_*n*−1_). Together, we can obtain the dynamics of cellular metabolism by optimizing growth rate *λ* in FBA, subject to constraining the PHB synthesis flux *ν*_*PHB*_(*t*_*n*_).

Using the extended dCAFBA model, we first examined the kinetic responses of growth rate *λ* and protein fraction of PHB synthesis *ϕ*_*P*._when PHB synthesis genes are induced from 0 to a given value *χ*_*P*0_ (Figure 4c). The simulated results showed that cellular growth rate *λ* decreased significantly upon inducing PHB gene expression at a titrating time *λ*_*C*_ =15 h and *χ*_*P*0_ =0.08, accompanying by an increase in PHB biosynthetic enzymes *ϕ*_*P*_ and PHB synthesis flux *v*_*PHB*_. We further identified the effects of gene expression level on PHB production by varying induction level *χ*_*P*0_. Model predictions suggested that increasing the level of PHB genes expression *χ*_*P*0_ enhanced PHB production accumulated over 24 hours (Figure S7). However, the PHB production started to decrease when the gene expression level *χ*_*P*0_ was set too high, for example, *χ*_*P*0_ > 0.32. This is likely because growth rate decreased rapidly when *χ*_*P*0_ was very high, limiting the increase of PHB protein fraction *ϕ*_*P*_. Although these predictions remain to be validated experimentally, the *in silico* results indicate that our dCAFBA model is a promising tool for characterizing the flux distribution and gene expression levels to enhance the production of desired products.

## Discussion

Microbes often encounter the fluctuating growth conditions, both naturally and artificially. In response to these variations, bacterial cells must globally regulate gene expression and proteome allocation, leading to metabolic reconfiguration and vice versa. ^1,17^ The cross-regulation between proteome allocation and metabolic flux determines the growth adaption to changing environments. ^21^ Studying how cells adjust metabolic flux and proteome allocation during growth shifts is crucial for understanding growth adaptation dynamics. To address this, we proposed a dynamic proteome-constrained flux balance approach for *E*.*coli*, termed dCAFBA, which combines flux balance analysis with the global regulation of proteome allocation. The resulting dCAFBA model enables the prediction of growth kinetics, dynamic reallocation of proteomic resources, and kinetic responses of metabolic flux under various transitions, such as nutrient (carbon substrate, amino acids) shifts and transcriptional signal variations, including enzyme expression induction, without detailing molecular gene regulations (Figure 5).

**Figure 5.**
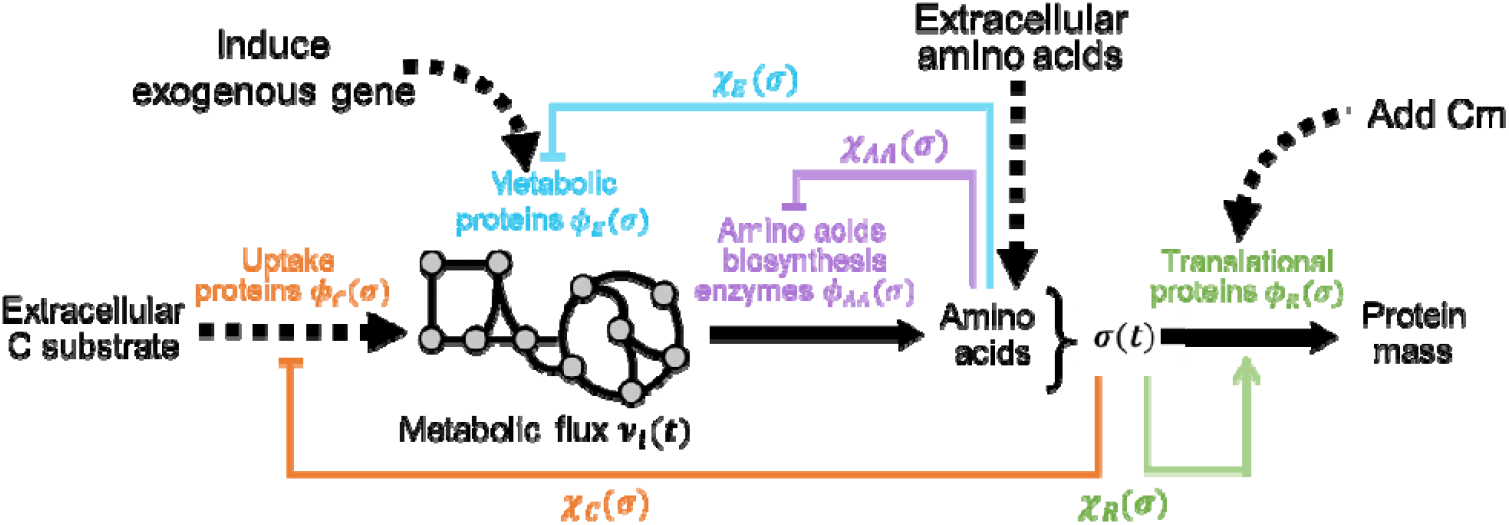
Summary of the dCAFBA model’s responses to various environmental fluctuations, including nutrient shifts (carbon and amino acids), regulatory signals to induce exogenous gene expression, and the addition of chloramphenicol.

The dCAFBA model successfully captured changes in growth rate and protein allocations during both carbon shifts (Figure S1) and amino acid down-shifts (Figure S3) without any adjustable parameters. Regarding flux distribution, systematic changes were observed during nutrient shifts. During the carbon up-shift from succinate to a combination of succinate and glycerol, the reaction catalyzed by fructose bisphosphatase (FBP) was expected to be active during the transition, as succinate and glycerol, being gluconeogenic carbon sources, require fluxes through glycolysis in reverse. Interestingly, we identified that fluxes carried by FBP and fructose bisphosphatase aldolase (FBA) dropped to zero immediately after the shift (Figure 2b). This prediction aligns with previous studies suggesting that the presence of fructose-1,6-biphosphate can suppress the glycerol uptake through feedback inhibition. ^23,24^ In the case of carbon down-shift, our model predicted an overshoot in the instantaneous growth rate of post-shift. Similar results have been found in other experimental and *in silico* studies, indicating that the growth rate overshoot results from a limitation switch from carbon uptake proteins to metabolic enzyme proteins. ^2,21^ The growth rate, being a direct readout of flux distribution through the metabolic network, implies that the overshoot phenomenon should also be observed in the kinetics of reaction fluxes, such as pyruvate dehydrogenase and citrate synthase (Figure 2c).

In the absence of amino acids, *de novo* AA biosynthesis is needed for sustaining cell growth. According to the end-product regulation, ^30^ AAB enzymes are not expressed simultaneously but hierarchically. Thus, cells preferentially allocate protein synthesis flux to the growth-limiting AAB enzyme group. By calculating the trigger time of each AAB enzyme group, we obtained the sequence of AAB enzymes express, consistent with the reported experimental observations (Figure 3b). ^3^ We also repeated simulations for other AA down-shift scenarios, such as the transition from glucose supplemented with casamino acids (cAA) to glucose minimal medium (Figure S5). ^4^ Interestingly, we found that met group, with the highest abundance in the amino acid-free minimal medium, was always the first to express in all simulated AA down-shift cases (the smallest trigger time), implying that met is the most bottlenecked enzyme group right after the shift. This observation is consistent with Björkeroth et al. ^31^ who reported that the met enzyme is synthesized first in the eukaryotic organism yeast. This is reasonable because methionine plays a crucial role in one-carbon metabolism, where sulfur from methionine is incorporated into cysteine (Figure S8). Due to the sequential express of AAB group enzymes, a growth lag was observed before growth recovery post-AA down-shift. ^3,4^ It has been shown that the growth lag can be varied by titrating the pre-shift expression of limiting enzymes. ^1^ The performance of dCAFBA also holds for *i*ML1515 model, highlighting our model predictions are not sensitive to the employed metabolic network (Figure S2, S6).

Traditionally, chemical signals are often used to activate the expression of metabolic enzymes, upregulating flux toward desired pathways in metabolic engineering applications. ^32,33^ In this study, we mimicked cellular dynamic responses perturbed by titrating genes expression for PHB production in *E*.*coli* using our dCAFBA model. The results revealed that an increase of PHB leads to a reduction in growth rate and disrupted metabolic states due to limited resources in a cell. As an intracellular product, PHB accumulation is always coupled with cellular growth, suggesting that any engineering strategy must consider its effects on growth rate and the reciprocal effect of growth on product synthesis due to the coupling between growth rate and proteomic and metabolic regulation. ^17,34^ Overall, the dCAFBA model proposed in our study provides a promising tool for predicting and analyzing cellular growth dynamics and metabolic response in changing environment. In addition to nutrient (carbon and amino acids) shifts and exogenous regulatory signals discussed here, the dCAFBA framework can, in principle, be extended to other perturbations, such as adding chloramphenicol, ^35^ when input such as regulatory functions and steady-state protein fractions are provided.

## STAR Methods

### Resource availability

#### Lead contact

Further information and requests for resources should be directed to and will be fulfilled by the lead contact, Xiongfei Fu.

## Materials availability

This study did not generate new materials.

## Data and code availability

Data supporting the findings of this study are available within the main text and Supplementary Information. The Custom-made simulation code is available at github: https://github.com/BaiYangBqdq/dCAFBA.

## Method details

### Model formulation and implementation

#### Model formulation

The dCAFBA method integrated ordinary differential equations (ODEs) with Flux Balance Analysis (FBA). The dynamic ODEs were solved with central differential method with a certain time step. FBA simulations were performed using COBRA toolbox executable in MATLAB (The MathWorks, version R2018b). ^36^ The GLPK Optimizer solver in combination with the toolbox, was used to solve the linear programming problems. In this study, the biomass equation was used to obtain the optimal solution of the metabolic model as described elsewhere. ^6^ Mathematically, the dCAFBA model can be described as follows:

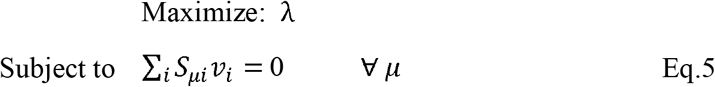

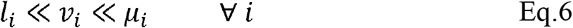

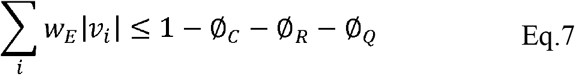

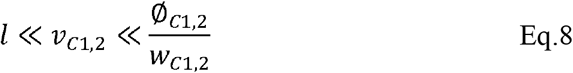

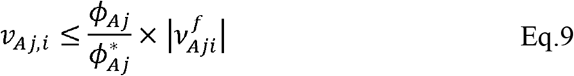

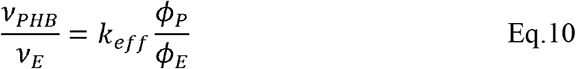

where *λ* is the flux that the biomass reaction carries, representing the growth rate. *S*_*ui*_ is the stoichiometric matrix, *u* is a vector of all metabolites, and *i* represents reactions. *l*_*i*_ and *u*_*i*_ represent the lower and upper bounds for the flux of each reaction *ν*_*i*_, respectively. The sum of all absolute metabolic fluxes *v*_*E*_ is constrained by: 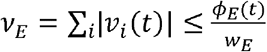 (Eq.7). Eq.8 is the flux boundary set on carbon influx during the carbon shift. Upon the AA down-shift, we incorporated Eq.9 to FBA as additional constraints on amino acid biosynthetic reactions. Eq.10 represents the specific constraints set on the PHB biosynthetic reaction upon the genetic perturbations, where *v*_*PHB*_ and *v*_*E*_ represent the fluxes going into PHB biosynthetic pathway and the sum of all metabolic fluxes, respectively. The parameter *k*_*eff*_ represents the competition for shared precursors between PHB synthesis and other metabolic reactions in the metabolic network. *w*_*C*_ and *w*_*E*_ characterize the cost of carbon uptake and metabolic proteins needed to carry one unit flux, determined by fitting the steady-state growth rates and protein partitions.

To model the dynamics of protein allocation, we divide the entire cellular proteome into four coarse-grained functional sectors according to their functions and their dependency on translational activity *σ* ^2^: (i) carbon-scavenging proteins (C-sector) including proteins devoted to carbon import and transport from environment, its protein fraction *ϕ*_*C*_ decreases with *σ*; (ii) translational proteins (R-sector) composed of ribosomal and affiliated proteins forming translational machinery, its protein fraction *ϕ*_*R*_ increases with *σ*; (iii) house-keeping proteins (Q-sector), its protein allocation *ϕ*_*Q*_ remains unchanged with *σ* ; (iv) metabolic enzymes (E-sector), consisting of catalytic enzymes related to metabolic pathways, its protein allocation *ϕ*_*E*_ exhibited complicated dependency on “. Given that each protein group fall into one of these four sectors, the *ϕ*_*E*_ can be calculated by*ϕ*_*E*_ = 1 − *ϕ*_*C*_ − *ϕ*_*R*_ − *ϕ*_*Q*_.

dCAFBA model is an integration of ODEs and FBA. At each time step, protein sectors *ϕ*_*R*_, *ϕ*_*C*1,2_, *ϕ*_*A,j*_, and *ϕ*_*P*_ are updated following a standard Euler scheme. The protein fractions *ϕ* follow a systems of ODEs (listed in Table S1). With the regulatory functions *χ*_*R*_, *χ*_*C*_, *χ*_*A,j*_ and *χ*_*P*_, we linearize this system into the following forward Euler scheme. We use these linearized equations to update the protein fractions after the solution to each time step has been computed. Then the metabolic enzymes *ϕ*_*E*_ can be computed at a given time point. With the calculated *ϕ*_*C*_, *ϕ*_*A,j*_, *ϕ*_*P*_, and *ϕ*_*E*_, we can further set the constraints on the corresponding reaction flux under different growth conditions by Eq. 7-10. Then, we compute the flux distribution *ν*_*i*_ by optimizing the biomass synthesis flux *λ*. We calculated the total amino acid synthesis flux (*ν*_*aa*_) to further determine the protein reallocation kinetics. Therefore, this iterative framework enabled us to calculate the flux redistribution processes. More details of the model implementation are described below.

#### Implementation of the carbon shift and transcriptional variation

##### Pre-shift growth

Since dCAFBA is an iterative method, it is necessary to set an initial value from which the dynamic analysis will integrate over time. The cell grows exponentially before the shift or titration (*t*< 0 *or t* < *t*_*C*_), thus, the initial value of growth rate *λ*: was set to the pre-shift growth rate which is experimentally measured. The initial value of ribosomal protein *ϕ*_*R*_ was set based on 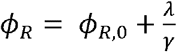. While the initial value of translational activity *σ* was set using 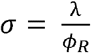. The initial value of *ϕ*_*C*_ was set based on 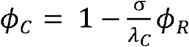. During the exponential growth, regulation functions (*χ*_*C*_, *χ*_*R*_, *χ*_*P*_) have the same value as their corresponding protein fraction (*ϕ*_*C*_, *ϕ*_*R*_, *ϕ*_*P*_), according to the kinetic equations of protein synthesis: 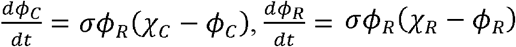 and 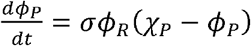.

The specific regulation functions *h*_1,2_ are introduced by a step function, based on the presence or absence of the cognate carbon substrate. This parameter *h*_1,2_ denotes the partition of a carbon substrate in a pair of co-utilized carbon sources in the case of carbon shift and are fitted to ensure the predicted growth rate and protein fractions matched experimental data. Then we can calculate the protein fractions for each substrate during the carbon shift by 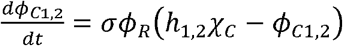. The metabolic enzymes *ϕ*_*E*_ can be calculated based on *ϕ*_*E*_ = 1 − *ϕ*_*R*_ − *ϕ*_*C*_ − *ϕ*_*Q*_. With the known *ϕ*_*E*_, *ϕ*_*C*1,2_ and *ϕ*_*P*_, we can bound the reaction fluxes based on Eq. 7, 8 and 10, respectively.

With the initial values of the above variables *ϕ*_*C*1,2_, *ϕ*_*R*_, *χ*_*C*_, *χ*_*R*_, *σ, ϕ*_*P*_, we can update the protein fractions to the next time point using a simple forward Euler method, with the time step Δ*t*_1_ = 0.01 *h*, to numerically solve the differential equations governing the protein dynamics.

With updated the values of *ϕ*_*C*1,2_, *ϕ*_*R*_ and *ϕ*_*P*_ at *t* + Δ*t*_1_, we then updated *v*_*aa*_(*t* + Δ*t*_1_), *λ*(*t* + Δ*t*_1_), *σ* (*t*+ Δ*t*_1_), *χ*_*R*_(*t* + Δ*t*_1_), and *χ*_*C*_ (*t* + Δ*t*_1_), following 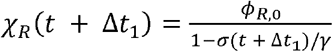, and 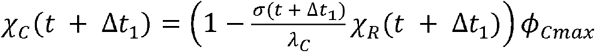. The regulation function *χ*_*P*_ was set to be 0 in the pre-shift as shown in Eq.4. *λ*_*C*_, *γ, ϕ*_*Cmax*_, and *ϕ*_*R*,0_ are constants determined by steady-state protein allocations under different growth conditions (supplemental tables).

##### At the instant of shift

At the instant of the shift (*t* = 0 or *t* = *t*_*C*_), all protein fractions must remain continuous. While, regulation functions and fluxes, however, can abruptly change at the moment of shift, depending on the initial condition *σ*(0). Translational activity was calculated by pre-shift growth rate and ribosome content using 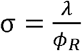. The change of *σ* at (*t* < 0) leads to changes in the values of the regulatory functions *χ*_*R*_ and *χ*_*C*_ using 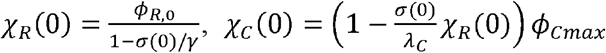. In the case of titrating PHB gene expression, *χ*_*P*_ was set as *χ*_*P0*_ at the instant of titration (*t*_*C*_). With updated the values of *χ*_*R*_ and *χ*_*C*_, we update the value of protein fractions *ϕ*_*C*1,2_, *ϕ*_*R*_, *ϕ*_*P*_, and *ϕ*_*E*_, which are used to further set the constraints on reaction flux.

##### Post-shift growth

Following the same procedure as described above for the pre-shift, we update the values of *ϕ*_C1,2_, *ϕ*_*R*_, *ϕ*_*P*_, *v*_*aa*_, *λ, σ, χ*_*R*_, and *χ*_*C*_ until all the variables reached a steady state.

#### Implementation of the AA down-shift

##### Pre-shift growth

In the pre-shift (*t* < 0), cells were growing exponentially in rich medium with amino acids. The pre-shift growth rate *λ* and protein fractions (including *ϕ*_*A,j*_ and *ϕ*_*R*_) were experimentaly determined, given in Table S3 and S6. During the steady state growth, regulation functions (*χ*_*R*_, *χ*_*A,j*_) have the same value as their corresponding protein abundance (*ϕ*_*R*_, *ϕ*_*A,j*_), according to the kinetic equations of protein synthesis: 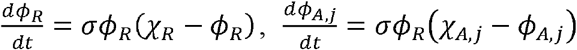. Thus, *χ*_*R*_(*t* < 0) = *ϕ*_*R*_(*t* < 0), *χ*_*A,j*_(*t* < 0)= *ϕ*_*A,j*_(*t* < 0) in the pre-shift steady state. With the values of growth rate *λ* (*t* < 0) and ribosomal fraction *ϕ*_*R*_(*t*< 0), translational activity was calculated by pre-shift growth rate and ribosome content using 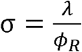. Because the carbon source is the same during the AA down-shift, we assumed the carbon uptake proteins *ϕ*_*C*_ to be unchanged, set as 3%. Referring to the previous study, ^17^ the house-keeping protein *ϕ*_*Q*_ was set as 0.45. With the values of *ϕ*_*R*_ (*t* < 0), *ϕ*_*C*_ (*t* < 0) and *ϕ*_*Q*_ given above, the *ϕ*_*E*_ can be calculated by *ϕ*_*E*_ (*t* < 0) = 1 − *ϕ*_*C*_ (*t* < 0) − *ϕ*_*R*_ (*t* < 0) − *ϕ*_*Q*_, which was then used for setting the constraints on the sum of all absolute metabolic fluxes *v*_*E*_ using equation when performing FBA.

##### At the instant of shift

Since, all protein fractions must be continuous at the instant of the shift (*t* = 0), we have *ϕ*_*R*_(0) = *ϕ*_*R*_ (*t* < 0), *ϕ*_*A,j*_ (*t* < 0) *ϕ*_*A,j*_ (*t* < 0). The initial values for fractional enzyme reserve *q*_*j*_(0) are thus obtained using 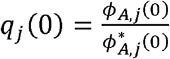. According to *q*_*j*_(0), we sorted AAB enzyme groups. The protein synthesis flux is allocated to the group with smallest *q*_*j*_(0) (the limiting group) using Eq. 3.

The removal of amino acids in the medium at *t* = 0 lead to the abruptly change in growth rate. Similarly, the translational activity also changes immediately from its value before the shift, according to 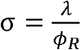. In the meanwhile, the change in *σ* at the instant of shift (*t* = 0) leads to changes in the values of the regulatory functions *χ*_*R*_ and *χ*_*A,tot*_ using 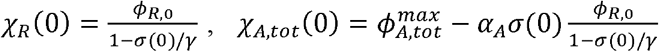. The parameters *α*_*A*_, *γ*, 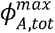, *ϕ*_*R*,0_ are constants determined from steady-state measurements (Table S3, S6). Finally, the regulatory function *χ*_*A,j*_(0) for each AAB enzyme group *j* was calculated based on *χ*^*A*^(0)*χ*^*A,tot*^(0). The relative allocation function *η*^*j*^ describes the portion of total AAB synthesis allocated to the enzyme group *j*. In the case of *q*_1_ (0) < *q*_2_ (0) < … < *q*_14_(0), we have *η*^1^ (0) = 1 and *η*^*j*^(0) (j ≠ 1) = 0. In the simulation of our AA downshift, the methionine group has the smallest *q*(0). With the *ϕ*_*E*_(0) (*ϕ*_*E*_(0) = 1 − *ϕ*_*C*_ − *ϕ*_*R*_(0) − *ϕ*_*Q*_) and *ϕ*_*A,j*_(0), we further constrained the reaction fluxes using Eq. 7 and Eq. 9, respectively.

##### Post-shift growth

Using updated regulatory functions at the instant of shift (*t* = 0), we can update the protein fractions to the next time point with the time step Δ*t*_2_ = 0.005 *h*, to numerically solve the differential equations that govern the protein dynamics.

With updated the values of *ϕ*_*R*_ and *ϕ*_*A,j*_ at *t* +Δ*t*_2_, we then updated *q*_*j*_ (*t* + Δ*t*_2_), *v*_*j*_ (*t* + Δ*t*_2_), *λ*(*t* + Δ*t*_2_), *σ*(*t* + Δ*t*_2_), *χ*_*R*_(*t* + Δ*t*_2_),*χ*_*A,tot*_(*t* + Δ*t*_2_), *η*_*j*_(*t* + Δ*t*_2_), and *χ*_*A,j*_ (*t* + Δ*t*_2_), following the same procedure as described above for updating their value at the instant shift (*t* = 0). The dynamics equations were simulated until all the variables reached a steady state.

## Acknowledgement

This work is partially supported by the National Key Research and Development Program of China (2023YFA0913900), the Strategic Priority Research Program of the Chinese Academy of Sciences (XDB0480000), NSFC (32071417), Guangdong Basic and Applied Basic Research Foundation (2022A1515110540), Shenzhen Science and Technology Program (ZDSYS20220606100606013).

## Author contributions

H.Y., Y.B. and X.F. conceived and designed the work. H.Y. carried out all simulations and drafted the manuscript. All the authors revised the manuscript.

## References

1. Basan, M., Honda, T., Christodoulou, D., Horl, M., Chang, Y.F., Leoncini, E., Mukherjee, A., Okano, H., Taylor, B.R., Silverman, J.M., et al. (2020). A universal trade-off between growth and lag in fluctuating environments. Nature 584, 470–474. 10.1038/s41586-020-2505-4.

2. Erickson, D.W., Schink, S.J., Patsalo, V., Williamson, J.R., Gerland, U., and Hwa, T. (2017). A global resource allocation strategy governs growth transition kinetics of Escherichia coli. Nature 551, 119-+.

3. Wu, C., Mori, M., Abele, M., Banaei-Esfahani, A., Zhang, Z., Okano, H., Aebersold, R., Ludwig, C., and Hwa, T. (2023). Enzyme expression kinetics by Escherichia coli during transition from rich to minimal media depends on proteome reserves. Nat Microbiol 8, 347–359. 10.1038/s41564-022-01310-w.

4. Zhu, M., and Dai, X. (2023). Stringent response ensures the timely adaptation of bacterial growth to nutrient downshift. Nat Commun 14, 467. 10.1038/s41467-023-36254-0.

5. Zhu, J., Chu, P., and Fu, X. (2023). Unbalanced response to growth variations reshapes the cell fate decision landscape. Nat Chem Biol 19, 1097–1104. 10.1038/s41589-023-01302-9.

6. Orth, J.D., Thiele, I., and Palsson, B.O. (2010). What is flux balance analysis? Nat Biotechnol 28, 245–248. 10.1038/nbt.1614.

7. McCloskey, D., Palsson, B.O., and Feist, A.M. (2013). Basic and applied uses of genome-scale metabolic network reconstructions of Escherichia coli. Mol Syst Biol 9, 661. 10.1038/msb.2013.18.

8. Yuan, H., Cheung, C.Y., Poolman, M.G., Hilbers, P.A., and van Riel, N.A. (2016). A genome-scale metabolic network reconstruction of tomato (Solanum lycopersicum L.) and its application to photorespiratory metabolism. Plant J 85, 289–304. 10.1111/tpj.13075.

9. Mahadevan, R., Edwards, J.S., and Doyle, F.J. (2002). Dynamic flux balance analysis of diauxic growth in Escherichia coli. Biophys J 83, 1331–1340.

10. Salvy, P., and Hatzimanikatis, V. (2021). Emergence of diauxie as an optimal growth strategy under resource allocation constraints in cellular metabolism. P Natl Acad Sci USA 118.

11. Waldherr, S., Oyarzun, D.A., and Bockmayr, A. (2015). Dynamic optimization of metabolic networks coupled with gene expression. J Theor Biol 365, 469–485. 10.1016/j.jtbi.2014.10.035.

12. Yang, L., Ebrahim, A., Lloyd, C.J., Saunders, M.A., and Palsson, B.O. (2019). DynamicME: dynamic simulation and refinement of integrated models of metabolism and protein expression. Bmc Syst Biol 13.

13. Chen, Y., and Nielsen, J.J.C.O.i.S.B. (2021). Mathematical modeling of proteome constraints within metabolism. 25, 50–56.

14. Mori, M., Hwa, T., Martin, O.C., De Martino, A., and Marinari, E. (2016). Constrained Allocation Flux Balance Analysis. Plos Comput Biol 12.

15. Zeng, H., and Yang, A. (2019). Quantification of proteomic and metabolic burdens predicts growth retardation and overflow metabolism in recombinant Escherichia coli. Biotechnol Bioeng 116, 1484–1495. 10.1002/bit.26943.

16. Scott, M., Gunderson, C.W., Mateescu, E.M., Zhang, Z.G., and Hwa, T. (2010). Interdependence of Cell Growth and Gene Expression: Origins and Consequences. Science 330, 1099–1102.

17. You, C.H., Okano, H., Hui, S., Zhang, Z.G., Kim, M., Gunderson, C.W., Wang, Y.P., Lenz, P., Yan, D.L., and Hwa, T. (2013). Coordination of bacterial proteome with metabolism by cyclic AMP signalling. Nature 500, 301–306.

18. Hui, S., Silverman, J.M., Chen, S.S., Erickson, D.W., Basan, M., Wang, J.L., Hwa, T., and Williamson, J.R. (2015). Quantitative proteomic analysis reveals a simple strategy of global resource allocation in bacteria. Mol Syst Biol 11.

19. Liao, C., Blanchard, A.E., and Lu, T. (2017). An integrative circuit-host modelling framework for predicting synthetic gene network behaviours. Nat Microbiol 2, 1658–1666. 10.1038/s41564-017-0022-5.

20. Molenaar, D., van Berlo, R., de Ridder, D., and Teusink, B. (2009). Shifts in growth strategies reflect tradeoffs in cellular economics. Mol Syst Biol 5, 323. 10.1038/msb.2009.82.

21. Yuan, H., Bai, Y., Li, X., and Fu, X.J.M.E. (2024). Cross-regulation between proteome reallocation and metabolic flux redistribution governs bacterial growth transition kinetics. 82, 60–68.

22. Bi, S., Kargeti, M., Colin, R., Farke, N., Link, H., and Sourjik, V.J.N.C. (2023). Dynamic fluctuations in a bacterial metabolic network. 14, 2173.

23. Hermsen, R., Okano, H., You, C., Werner, N., and Hwa, T. (2015). A growth-rate composition formula for the growth of E.coli on co-utilized carbon substrates. Mol Syst Biol 11, 801. 10.15252/msb.20145537.

24. Zwaig, N., and Lin, E.C. (1966). Feedback inhibition of glycerol kinase, a catabolic enzyme in Escherichia coli. Science 153, 755–757. 10.1126/science.153.3737.755.

25. Reed, J.L., Vo, T.D., Schilling, C.H., and Palsson, B.O. (2003). An expanded genome-scale model of Escherichia coli K-12 (iJR904 GSM/GPR). Genome Biology 4.

26. Monk, J.M., Lloyd, C.J., Brunk, E., Mih, N., Sastry, A., King, Z., Takeuchi, R., Nomura, W., Zhang, Z., and Mori, H.J.N.b. (2017). i ML1515, a knowledgebase that computes Escherichia coli traits. 35, 904–908.

27. King, Z.A., Drager, A., Ebrahim, A., Sonnenschein, N., Lewis, N.E., and Palsson, B.O. (2015). Escher: A Web Application for Building, Sharing, and Embedding Data-Rich Visualizations of Biological Pathways. Plos Comput Biol 11, e1004321. 10.1371/journal.pcbi.1004321.

28. McAdam, B., Brennan Fournet, M., McDonald, P., and Mojicevic, M. (2020). Production of Polyhydroxybutyrate (PHB) and Factors Impacting Its Chemical and Mechanical Characteristics. Polymers (Basel) 12. 10.3390/polym12122908.

29. Zheng, Y., Yuan, Q., Yang, X., and Ma, H. (2017). Engineering Escherichia coli for poly-(3-hydroxybutyrate) production guided by genome-scale metabolic network analysis. Enzyme Microb Technol 106, 60–66. 10.1016/j.enzmictec.2017.07.003.

30. Elf, J., Berg, O.G., and Ehrenberg, M. (2001). Comparison of repressor and transcriptional attenuator systems for control of amino acid biosynthetic operons. J Mol Biol 313, 941–954. 10.1006/jmbi.2001.5096.

31. Bjorkeroth, J., Campbell, K., Malina, C., Yu, R., Di Bartolomeo, F., and Nielsen, J. (2020). Proteome reallocation from amino acid biosynthesis to ribosomes enables yeast to grow faster in rich media. Proc Natl Acad Sci U S A 117, 21804–21812. 10.1073/pnas.1921890117.

32. Hartline, C.J., Schmitz, A.C., Han, Y., and Zhang, F. (2021). Dynamic control in metabolic engineering: Theories, tools, and applications. Metab Eng 63, 126–140. 10.1016/j.ymben.2020.08.015.

33. Burg, J.M., Cooper, C.B., Ye, Z., Reed, B.R., Moreb, E.A., and Lynch, M.D.J.C.O.i.C.E. (2016). Large-scale bioprocess competitiveness: the potential of dynamic metabolic control in two-stage fermentations. 14, 121–136.

34. Klumpp, S., Zhang, Z., and Hwa, T. (2009). Growth rate-dependent global effects on gene expression in bacteria. Cell 139, 1366–1375. 10.1016/j.cell.2009.12.001.

35. Wu, C. (2022). Proteomic allocation by E. coli during growth transitions, from strategy to mechanism (University of California, San Diego).

36. Heirendt, L., Arreckx, S., Pfau, T., Mendoza, S.N., Richelle, A., Heinken, A., Haraldsdottir, H.S., Wachowiak, J., Keating, S.M., Vlasov, V., et al. (2019). Creation and analysis of biochemical constraint-based models using the COBRA Toolbox v.3.0. Nat Protoc 14, 639–702. 10.1038/s41596-018-0098-2.

