## Supplemental Table S1-10, Supplemental Figure S1-8 for "A global resource constrained model to predict metabolic flux dynamics in fluctuating environments"

**Supplemental tables**

Table S1. Model variables used in this study. In this table, we summarized all the variables used in the model. Here the subscript $i$ represents a protein group. The subscript $j$ only represents AAB enzyme groups. The asterisk represents the value in the post-shift steady-state.

| **Variables** | **Related equation** | **Description** |
| --- | --- | --- |
| $\sigma$ | $\sigma\left( t \right)\equiv\frac{\lambda(t)}{\phi_{R}(t)}$ | translational activity |
| $\chi_{R}$ | $\chi_{R}(t)=\frac{\phi_{Rb,0}}{1-\sigma/\gamma}$ | regulation function of ribosomal protein |
| $\chi_{C}$ | $\chi_{C}\left( t \right)=\left( 1-\frac{\sigma}{\lambda_{C}}\chi_{R} \right)\phi_{Cmax}$ | regulation function of catabolic protein allocation |
| $\chi_{A,tot}$ | $\chi_{A,tot}\left( t \right)=\phi_{A,tot}^{max}-\alpha_{A}\sigma\frac{\phi_{Rb,0}}{1-\sigma/\gamma}$ | regulation function of total AAB enzymes |
| $\chi_{A,j}$ | $\chi_{A,j}\left( t \right)=\eta_{j}\left( t \right)\chi_{A,tot}(t)$ | regulation function of individual AAB enzyme group |
| $\eta_{j}$ | $\eta_{j}\left( t \right)=\left\{ \begin{aligned} 0 j>\tilde{j}(t) \\ \phi_{A,j}^{*}/\sum_{m=1}^{\tilde{j}(t)} \phi_{A,m}^{*} j\leq\tilde{j}(t) \end{aligned} \right.$ | fraction of protein synthesis for total AAB enzyme allocated to group *j* |
| $\tilde{j}(t)$ |  | the number of AAB enzyme groups being synthesized at time t |
| $q_{j}$ | $q_{j}(t)\equiv\frac{\phi_{A,j}(t)}{\phi_{A,j}^{*}}$ | abundance of AAB enzyme group *j* relative to its post-shift steady-state |
| $\chi_{P}$ | $\chi_{P}\left( t \right)= \left\{ \begin{aligned} 0 t<t_{C} \\ \chi_{P0} t\geq t_{C} \end{aligned} \right.$ | regulation function of PHB biosynthetic enzymes |
| $\sigma$ | $\frac{d\sigma\left( t \right)}{dt}=\beta\frac{1}{\phi_{R}\left( t \right)}\left( \frac{{dv}_{aa}}{dt}-v_{aa}\left( t \right)\sigma\left( t \right)\left( \chi_{C}-\chi_{R} \right) \right)$ | dynamics of $\sigma$ used in carbon shift or titrating PHB gene expression |
| $\phi_{R}$ | $\frac{d\phi_{R}}{dt}=\sigma\phi_{R}(\chi_{R}-\phi_{R})$ | mass fraction of ribosomal protein over total protein |
| $\phi_{C}$ | $\frac{d\phi_{C1,2}}{dt}=\sigma\phi_{R}\left( h_{1,2}\chi_{C}-\phi_{C} \right)$ | mass fraction of catabolic protein over total protein |
| $\phi_{A,tot}$ | $\phi_{A,tot}=\sum_{j} \phi_{A,j}$ | mass fraction of the total AAB enzymes over total protein |
| $\phi_{A,j}$ | $\frac{d\phi_{A,j}}{dt}=\sigma\phi_{R}(\chi_{A,j}-\phi_{A,j})$ | mass fraction of individual AAB enzyme group |
| $\phi_{E}$ | $\phi_{E}\left( t \right)=1-\phi_{C}\left( t \right)-\phi_{R}\left( t \right)-\phi_{Q}$ | mass fraction of metabolic protein over total protein |
| $\phi_{P}$ | $\frac{d\phi_{P}}{dt}=\sigma\phi_{R}\left( \chi_{P}-\phi_{P} \right)$ | mass fraction of PHB biosynthetic protein over total protein |
| $\nu_{aa}$ |  | protein synthesis flux |
| $\nu_{C}$ |  | carbon uptake rate |
| $\nu_{E}$ |  | the total metabolic flux |
| $\nu_{ji}$ |  | flux carried by each AAB biosynthetic reaction |
| $\nu_{PHB}$ |  | PHB biosynthesis flux |
| $\lambda$ |  | growth rate |

Table S2. Description of parameters.

| *Symbol* | Description |
| --- | --- |
| $\gamma$ | inverse slope of growth laws |
| $\lambda_{C}$ | strain specific constant of growth laws |
| $\phi_{R,0}$ | a growth rate-independent offset of R sector |
| $\phi_{Q}$ | fraction of house-keeping protein |
| $\phi_{C}^{max}$ | maximum of proteome attainable to C, E and R sectors |
| $\beta$ | conversion factor between moles and gram |
| $h_{1,2}$ | carbon-specific allocation |
| $\omega_{E}$ | enzyme cost per unit metabolic influx |
| $\omega_{C}$ | enzyme cost per unit carbon influx |
| $\phi_{A,tot}^{max}$ | maximum of AAB enzyme at a given condition |
| $\alpha_{A}$ | relation parameter between AAB enzyme and growth rate |

Table S3. The parameters and constants used in the simulation of carbon shifts using *E.coli i*JR904 metabolic model.

|  | Substarte1 | Substarte2 | $\lambda_{1}$  (h^-1^) | $\lambda_{12}$  (h^-1^) | $w_{C1}$ | $w_{C2}$ | $h_{1}$ | $h_{2}$ | $\phi_{C1}^{max}$ | $\phi_{C2}^{max}$ | $w_{E}$ | $\gamma$  (h^-1^) | $\lambda_{C}$  (h^-1^) | $\phi_{Q}$ | $\phi_{R,0}$ | $\beta$  (g/mmol) |
| --- | --- | --- | --- | --- | --- | --- | --- | --- | --- | --- | --- | --- | --- | --- | --- | --- |
|  |  |  |  |  | gh/mmol | |  |  |  |  | gh/mmol |  |  |  |  |  |
| Up-shift | pyruvate | glycerol | 0.60 | 0.86 | 9.05×10^-3^ | 4.07×10^-3^ | 1 | 0.35 | 0.35 | 0.34 | 8.3×10^-4^ | 11.02 | 1.17 | 0.45 | 0.049 | 0.1968 |
|  | succinate | glycerol | 0.46 | 0.78 | 1.96×10^-2^ | 8.87×10^-3^ | 1 | 0.25 | 0.30 | 0.42 | 8.3×10^-4^ | 11.02 | 1.17 | 0.45 | 0.049 | 0.1968 |
| Down-shift | pyruvate | glycerol | 0.68 | 0.96 | 5.05×10^-3^ | 3.07×10^-3^ | 0.6 | 1 | 0.47 | 0.41 | 8.3×10^-4^ | 11.02 | 1.17 | 0.45 | 0.049 | 0.1968 |

Value of $w_{E}$ is obtained from Mori et al. ^1^

Values of $\gamma$, $\lambda_{C}$, $\phi_{Q}$ and $\phi_{R,0}$ were obtained from Erickson et al. ^2^

Table S 4. The parameters and constants used in the simulation of carbon shifts using e *E.coli i*ML1515 metabolic model.

|  | Substarte1 | Substarte2 | $\lambda_{1}$  (h^-1^) | $\lambda_{12}$  (h^-1^) | $w_{C1}$ | $w_{C2}$ | $h_{1}$ | $h_{2}$ | $\phi_{C1}^{max}$ | $\phi_{C2}^{max}$ | $w_{E}$ | $\gamma$  (h^-1^) | $\lambda_{C}$  (h^-1^) | $\phi_{Q}$ | $\phi_{R,0}$ | $\beta$  (g/mmol) |
| --- | --- | --- | --- | --- | --- | --- | --- | --- | --- | --- | --- | --- | --- | --- | --- | --- |
|  |  |  |  |  | gh/mmol | |  |  |  |  | gh/mmol |  |  |  |  |  |
| Up-shift | succinate | glycerol | 0.46 | 0.78 | 1.86×10^-2^ | 4.87×10^-3^ | 1 | 0.25 | 0.35 | 0.40 | 3.3×10^-4^ | 11.02 | 1.17 | 0.45 | 0.049 | 0.1970 |
| Down-shift | pyruvate | glycerol | 0.68 | 0.96 | 4.35×10^-3^ | 1.0×10^-3^ | 0.70 | 1 | 0.45 | 0.48 | 4.2×10^-4^ | 11.02 | 1.17 | 0.45 | 0.049 | 0.1970 |

Table S5. The AAB enzyme groups used in the simulation of AA down-shift.

| **AAB enzyme group** | **Genes** | **Reactions ID in *i*JR904** | **Reactions ID in *i*ML1515** |
| --- | --- | --- | --- |
| arg group | argA, argB, argC, argD, argE, argF, argG, argH, argI | ACGS, ACGK, AGPR, ACOTA, ACODA, NACODA, ARGSS, ARGSL, OCBT | ACGS, ACGK, AGPR, ACOTA, ACODA, ARGSS, ARGSL, OCBT |
| aro group | aroA, aroB, aroC, aroD, aroE, aroF, aroG, aroH, aroK, aroL | PSCVT, DHQS, CHORS, DHQD, SHK3Dr, DDPA, SHKK | PSCVT, DHQS, CHORS, DHQTi, SHK3Dr, DDPA, SHKK |
| cys group | cysC, cysD, cysE, cysH, cysI, cysJ, cysK, cysM, cysN | ADSK, SADT2, SERAT, PAPSR, SULR, CYSS | ADSK, SADT2, SERAT, PAPSR, SULR, CYSS |
| glt group | gdhA, gltB, gltD | GLUDy, GLUSy | GLUDy, GLUSy |
| his group | hisA, hisB, hisC, hisD, hisF, hisG, hisH, hisI | PRMICIi, IGPDH, HISTP, HSTPT, HISTD, IG3PS, ATPPRT, PARTPP, PRAMPC | PRMICI, IGPDH, HISTP, HSTPT, HISTD, IG3PS, ATPPRT, PARTPP, PRAMPC |
| ilv group | ilvA, ilvB, ilvC, ilvD, ilvE, ilvG_1, ilvH, ilvI, ilvM, ilvN | THRD_L, ACHBS, ACLS, KARA1i, KARA2i, DHAD1, DHAD2, ILETA, VALTA, ACHBS, ACLS | THRD_L, ACHBS, ACLS, KARA1, KARA2, DHAD1, DHAD2, ILETA, VALTA, ACHBS, ACLS |
| leu group | leuA, leuB, leuC, leuD | IPPS, IMPD, OMCDC, IPPMIa, IPPMIb | IPPS, IMPD, OMCDC, IPPMIa, IPPMIb |
| lys group | asd, dapA, dapB, dapD, dapE, dapF, lysA, lysC | ASAD, DHDPS, DHDPRy, THDPS, SDPDS, DAPE, DAPDC | ASAD, DHDPS, DHDPRy, THDPS, SDPDS, DAPE, DAPDC |
| met group | metA, metB, metC, metE, metH, metL | HSST, SHSL1, CYSTL, METS, ASPK, HSDy | HSST, SHSL1, CYSTL, METS, ASPK, HSDy |
| other group | alaA, alaC, asnA, asnB, aspC, glnA, glyA, proA, proB, tyrB, proC | ALATA_L, ASNS2, ASNS1, ASPTA, GLNS, THRAr, GHMT2, G5SD, GLU5K, P5CR | ALATA_L, ASNS2, ASNS1, ASPTA, GLNS, THRA2, GHMT2r, G5SD, GLU5K, P5CR |
| phetyr group | pheA, tyrA, tyrB | CHORM, PPNDH, PHETA1, LEUTAi, TYRTA | CHORM, PPNDH, PHETA1, LEUTAi, TYRTA |
| ser group | serA, serB, serC | PGCD, PSP_L, PSERT | PGCD, PSP_L, PSERT |
| thr group | thrA, thrB, thrC | HSK, THRS | HSK, THRS |
| trp group | trpA, trpB, trpC, trpD, trpE | TRPS1, TRPS2, TRPS3, PRAIi, IGPS, ANS, ANPRT | TRPS1, TRPS2, TRPS3, PRAIi, IGPS, ANS, ANPRT |

Table S6. Experimental data used in the simulation of AA down-shift, including the transition from glycerol supplemented 18 AA (i.e., glycerol +18AA) or 18 AA excluding serine (i.e., glycerol +18AA-Ser) to glycerol minimal medium (post-shift), and the transition from glucose supplemented cAA (i.e., glucose +cAA) to glucose minimal medium (post-shift).

|  | Pre-shift Glycerol+18AA**^1^** | Pre-shift  Glycerol+18AA-Ser**^1^** | Post-shift  Glycerol**^1^** | Pre-shift  Glucose+cAA**^2^** | Post-shift Glucos**e^2^** |
| --- | --- | --- | --- | --- | --- |
| $\phi_{A,met}$ | 0.60‰ | 1.05‰ | 32.2‰ | 1.22‰ | 57‰ |
| $\phi_{A,arg}$ | 0.14‰ | 0.20‰ | 4.71‰ | 0.559‰ | 8.32‰ |
| $\phi_{A,leu}$ | 0.25‰ | 0.51‰ | 5.85‰ | 0.47‰ | 5.74‰ |
| $\phi_{A,trp}$ | 0.12‰ | 0.16‰ | 2.31‰ | 2.65‰ | 6.58‰ |
| $\phi_{A,glt}$ | 0.72‰ | 1.12‰ | 7.23‰ | 0.94‰ | 8.88‰ |
| $\phi_{A,ilv}$ | 1.60‰ | 1.80‰ | 13.1‰ | 3.22‰ | 15.51‰ |
| $\phi_{A,his}$ | 0.74 ‰ | 1.38‰ | 4.15‰ | 0.94‰ | 3.63‰ |
| $\phi_{A,ser}$ | 1.00‰ | 1.25‰ | 3.79‰ | 1.0‰ | 7.71‰ |
| $\phi_{A,lys}$ | 2.63‰ | 2.75‰ | 6.32‰ | 2.83‰ | 8.69‰ |
| $\phi_{A,thr}$ | 2.40‰ | 1.21‰ | 4.13‰ | 1.35‰ | 5.67‰ |
| $\phi_{A,aro}$ | 1.97‰ | 1.67‰ | 2.97‰ | 1.98‰ | 4.33‰ |
| $\phi_{A,phetyr}$ | 0.65 ‰ | 0.63‰ | 1.09‰ | 0.41‰ | 1.05‰ |
| $\phi_{A,cys}$ | 8.66 3‰ | 6.51‰ | 9.88‰ | 1.1‰ | 11.48‰ |
| $\phi_{A,other}$ | 11.1‰ | 10.5‰ | 11.5‰ | 14.9‰ | 15.37‰ |
| $\phi_{A,tot}^{max}$ | 18.1% | 21.8% | nd | 0.322 | nd |
| $\alpha_{A}$ | 0.11 h | 0.16 h | nd | 0.182 h | nd |
| $\phi_{R,0}$ | 4.45% | | | | |
| $\gamma$ | 8.35/h | | | | |
| $\lambda$ | 1.40 h | 1.17 h | 0.68 h | 1.58 h | 0.95 h |

**^1^**Data were obtained from Wu et al. ^3^

**^2^**Data were obtained from Zhu & Dai ^4^

$\phi_{A,tot}^{max}$ and $\alpha_{A}$ were calculated based on pre-shift and post-shift growth rate and total AAB enzyme abundance as described in Wu et al. ^3^

Table S7. The constants of the substrates used in the simulation of AA down-shift with the *E.coli i*JR904 metabolic model, including the transition from glycerol supplemented 18 AA (i.e., glycerol +18AA) or 18 AA excluding serine (i.e., glycerol +18AA-Ser) to glycerol minimal medium, ^3^ and the transition from glucose supplemented cAA (i.e., glucose +cAA) to glucose minimal medium. ^4^ The unit of $w_{E}$ is gh/mmol.

|  | Pre-shift Glycerol+18AA | Pre-shift  Glycerol+18AA-Ser | Post-shift  Glycerol | Pre-shift  Glucose+cAA | Post-shift Glucose |
| --- | --- | --- | --- | --- | --- |
| $\phi_{Q}$ | 0.45 | 0.45 | 0.45 | 0.40 | 0.45 |
| $w_{E}$ | 2.0×10^-5^ | 3.0×10^-4^ | 8.3×10^-4^ | 1.0×10^-5^ | 8.3×10^-4^ |
| $\phi_{C}$ | 0.03 | 0.03 | 0.03 | 0.03 | 0.03 |

Table S8. The constants of the substrates used in the simulation of AA down-shift with the *E.coli i*ML1515 metabolic model: the transition from glycerol supplemented 18 AA (i.e., glycerol +18AA) to glycerol minimal medium. ^3^

| Items | Pre-shift Glycerol+18AA | Post-shift  Glycerol |
| --- | --- | --- |
| $\phi_{Q}$ | 0.35 | 0.35 |
| $w_{E}$ | 2.8×10^-4^ (gh/mmol) | 8.3×10^-4^ (gh/mmol) |
| $\phi_{C}$ | 0.0 | 0.0 |

Table S9. The parameters used in the simulation of PHB production by dCAFBA.

| Substrate | $\lambda$ (h^-1^) | $\omega_{C}$(gh/mmol) | $\omega_{E}$(gh/mmol) | $\phi_{Cmax}$ | $k_{eff}$ | $t_{C}$(h) |
| --- | --- | --- | --- | --- | --- | --- |
| glucose | 0.85 | 1.0×10^-3^ | 4.1×10^-4^ | 0.24 | 0.1 | 15 |

Table S10. Amino acids included in the studied medium.

Cysteine and tyrosine are excluded in 18AAs because of their instability (cysteine) and poor solubility (tyrosine).

| **Amino acids** | **18AA** | **18AA-Ser** | **cAA** |
| --- | --- | --- | --- |
| alanine | + | + | + |
| arginine | + | + | + |
| aspartate | + | + | + |
| asparagine | + | + | + |
| cysteine | - | - | + |
| glutamate | + | + | + |
| glutamine | + | + | + |
| glycine | + | + | + |
| histidine | + | + | + |
| isoleucine | + | + | + |
| leucine | + | + | + |
| lysine | + | + | + |
| methionine | + | + | + |
| phenylalanine | + | + | + |
| proline | + | + | + |
| serine | + | - | + |
| threonine | + | + | + |
| tryptophan | + | + | - |
| tyrosine | - | - | + |
| valine | + | + | + |

**Supplemental figures**

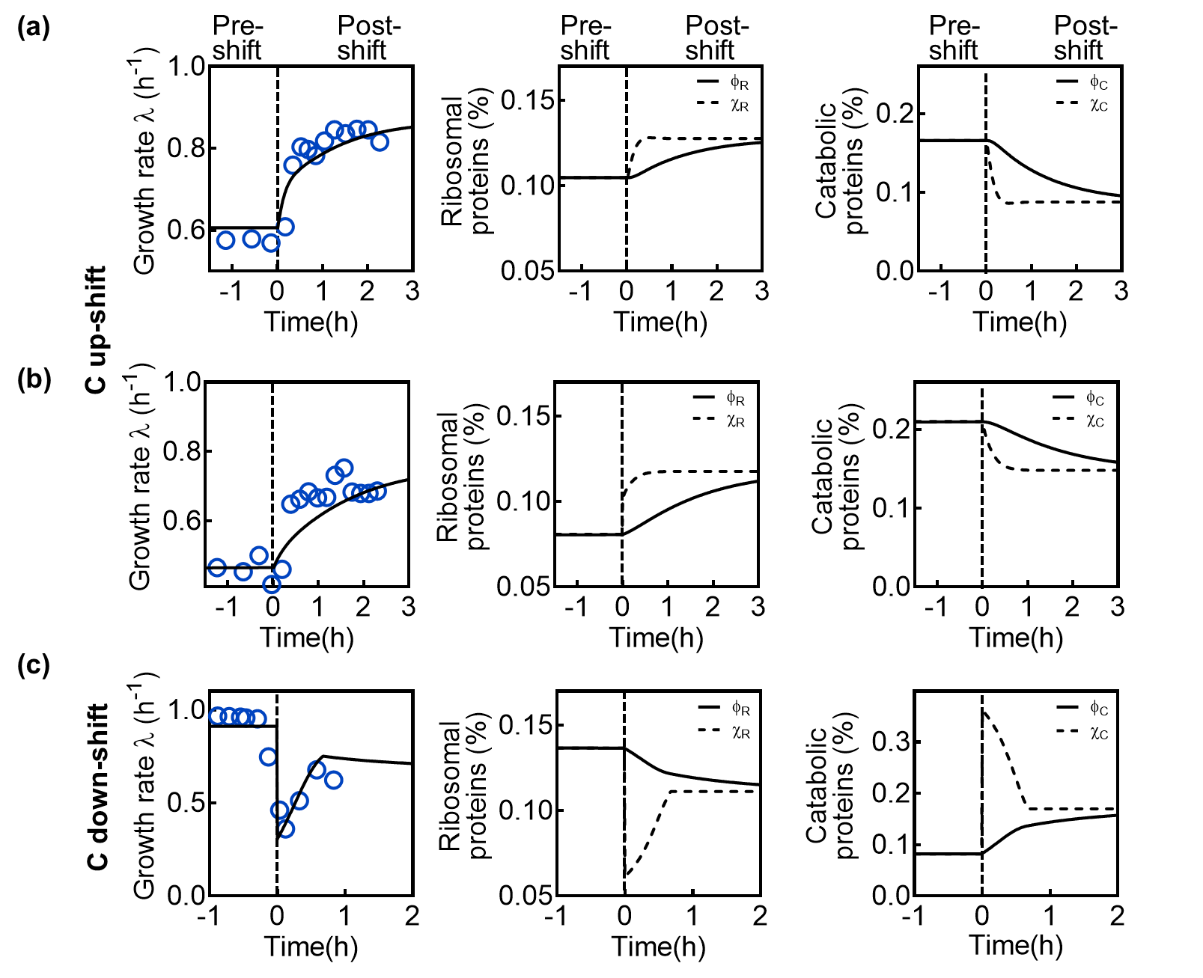

Figure S1. Dynamics of growth rate $\lambda\left( t \right)$, regulatory functions $\chi_{C}$ and $\chi_{R}$, protein fractions $\phi_{R}$ and $\phi_{C}$ in the conditions of transition from pyruvate to pyruvate and glycerol (a), from succinate to succinate and glycerol (b), and from pyruvate and glycerol to pyruvate (c). Blue circles are experimental data extracted from the referenced literature ^2^ using the online version of WebPlotDigitizer 4.2 (<https://automeris.io/WebPlotDigitizer/>).

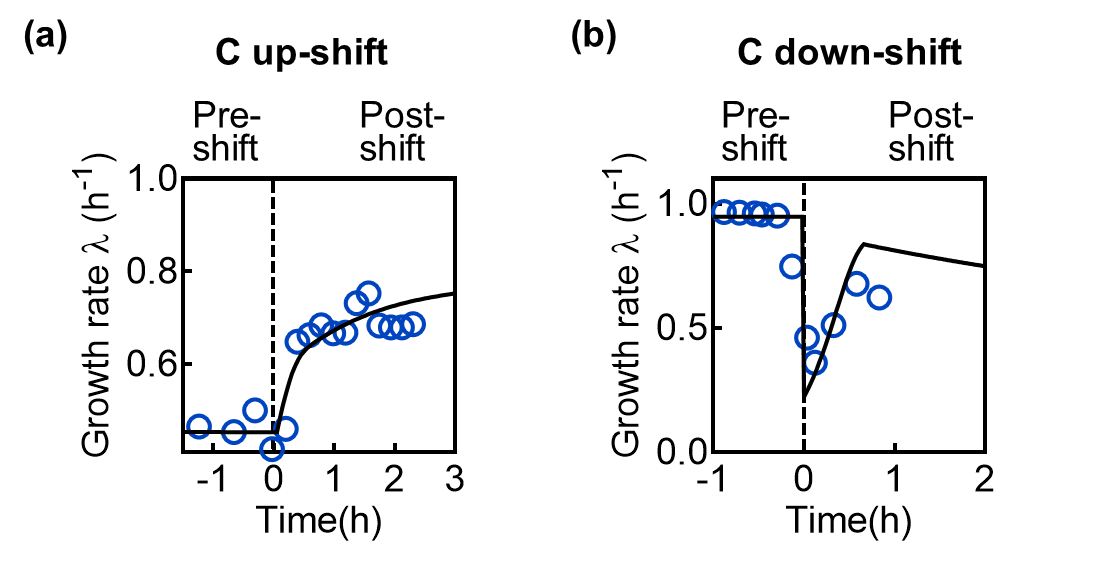

Figure S2. Dynamics of growth rate $\lambda\left( t \right)$ in the conditions of transition from succinate to succinate and glycerol (a), and from pyruvate and glycerol to pyruvate (b) using *i*ML1515 model. Blue circles are experimental data extracted from the referenced literature ^2^.

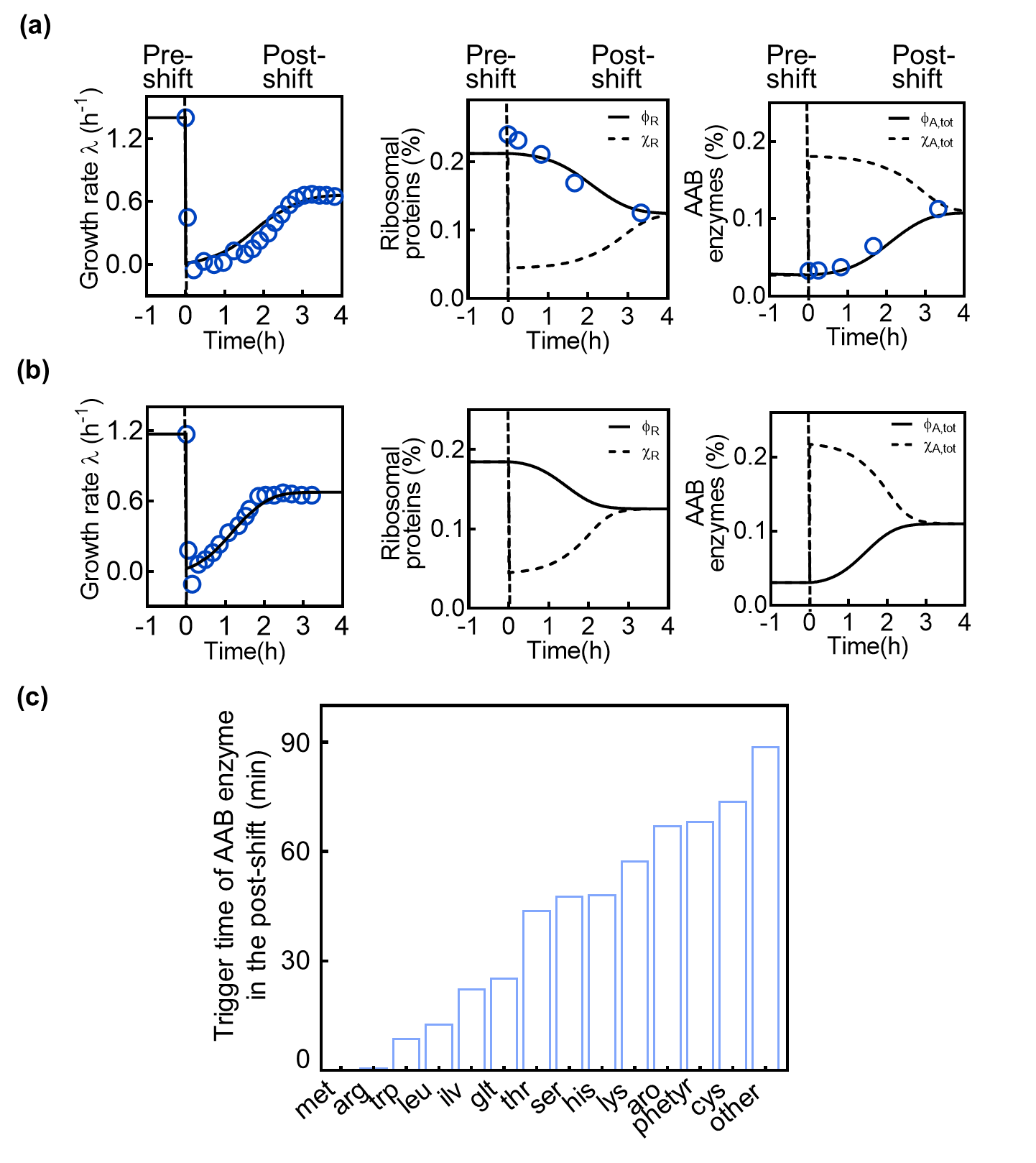

Figure S3. Dynamics of growth rate $\lambda\left( t \right)$, regulatory functions $\chi_{AA}$ and $\chi_{R}$, protein fractions $\phi_{R}$ and $\phi_{AA}$ in the conditions of transition from glycerol supplemented with 18AAs to glycerol (a), glycerol supplemented with 18AAs-serine to glycerol (b). Trigger time of each AAB enzymes in the post shift from glycerol with 18AAs to glycerol (c). Blue circles are experimental data from Wu et al. ^3^

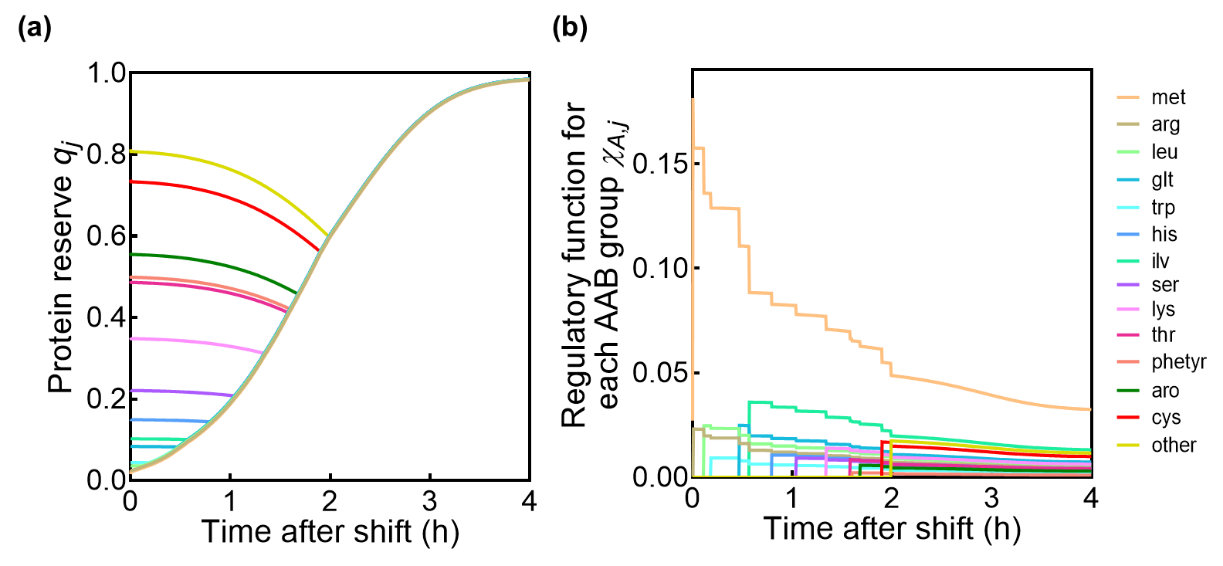

Figure S4. Protein reserve (a) and regulation function of each AAB (b) in the conditions of transition from glycerol supplemented with 18AA to glycerol.

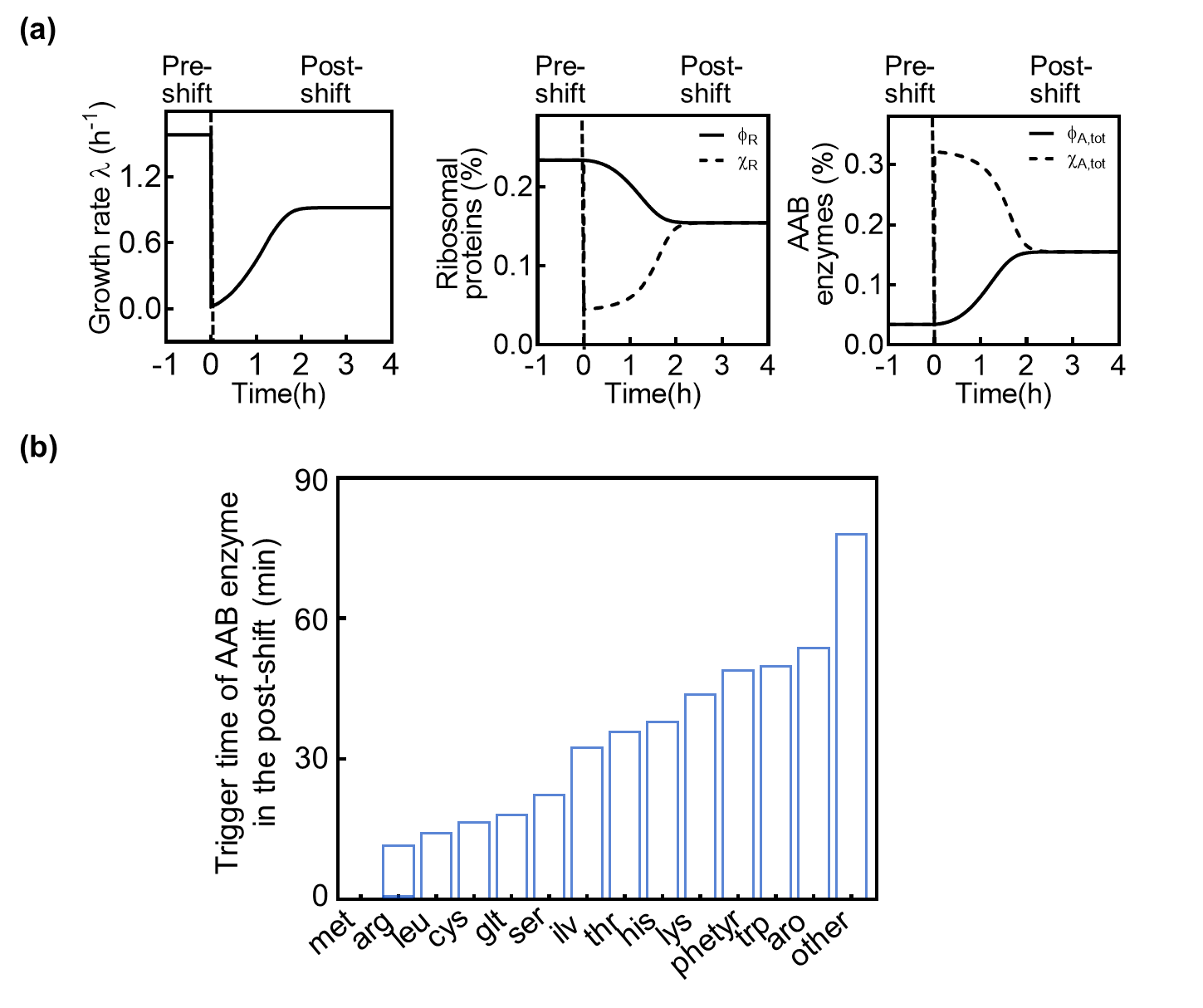

Figure S5. Dynamics of growth rate $\lambda\left( t \right)$, regulatory functions $\chi_{AA}$ and $\chi_{R}$, protein fractions $\phi_{R}$ and $\phi_{AA}$ in the conditions of transition from glucose with cAA to glucose (a, this case was studied in the work of Zhu and Dai, ^4^ and trigger time of each AAB enzymes in the post shift (b).

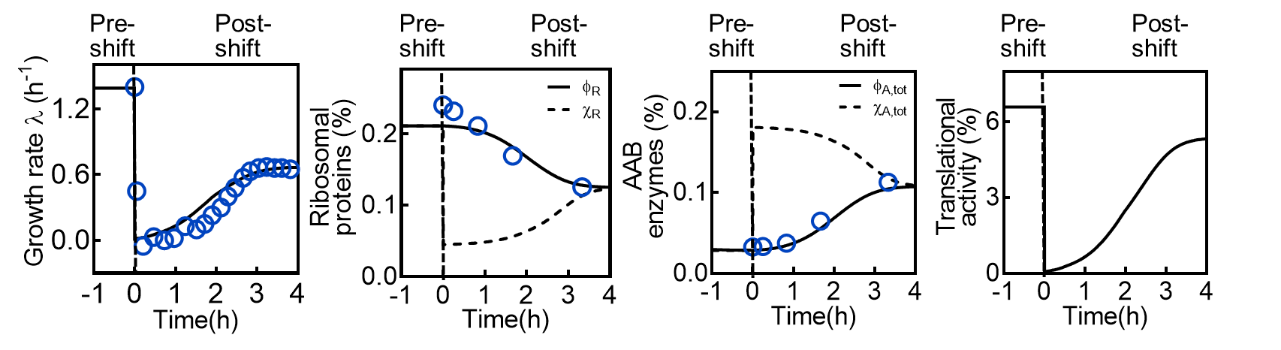

Figure S6. Dynamics of growth rate $\lambda\left( t \right)$, regulatory functions $\chi_{AA}$ and $\chi_{R}$, protein fractions $\phi_{R}$ and $\phi_{AA}$, and translational activity $\sigma$ during the transition from glycerol with 18AA to none using *i*ML1515 model. Blue circles are experimental data from Wu et al. ^3^

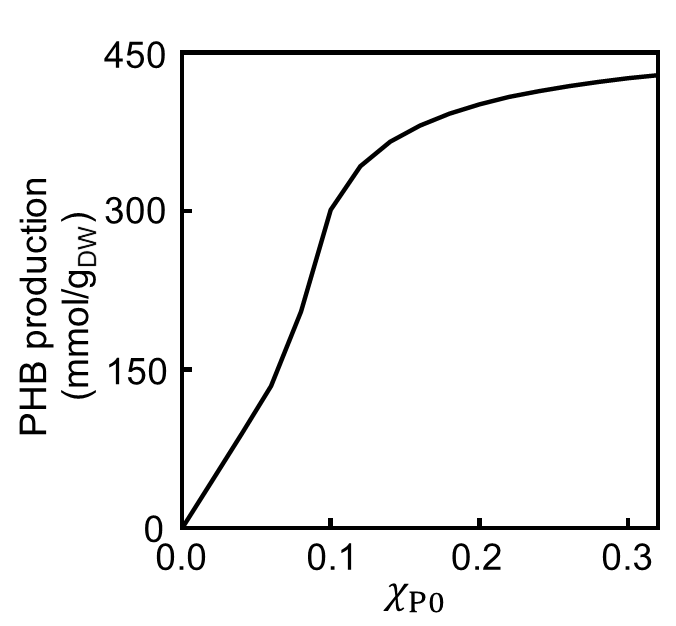

Figure S7. Changes of PHB production under varying $\chi_{P0}$.

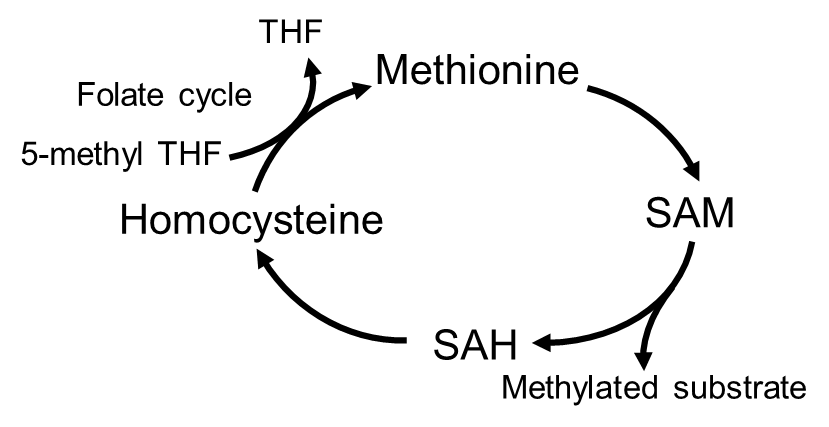

Figure S8. Methionine metabolism and its connections to one carbon metabolism. SAM, S-adenosyl methionine; SAH, S-adenosyl homocysteine; THF, tetrahydrofolate.

**Reference**

1.	Mori, M., Hwa, T., Martin, O.C., De Martino, A., and Marinari, E. (2016). Constrained Allocation Flux Balance Analysis. Plos Comput Biol *12*.

2.	Erickson, D.W., Schink, S.J., Patsalo, V., Williamson, J.R., Gerland, U., and Hwa, T. (2017). A global resource allocation strategy governs growth transition kinetics of Escherichia coli. Nature *551*, 119-+.

3.	Wu, C., Mori, M., Abele, M., Banaei-Esfahani, A., Zhang, Z., Okano, H., Aebersold, R., Ludwig, C., and Hwa, T. (2023). Enzyme expression kinetics by Escherichia coli during transition from rich to minimal media depends on proteome reserves. Nat Microbiol *8*, 347-359. 10.1038/s41564-022-01310-w.

4.	Zhu, M., and Dai, X. (2023). Stringent response ensures the timely adaptation of bacterial growth to nutrient downshift. Nat Commun *14*, 467. 10.1038/s41467-023-36254-0.
